# A gene drive does not spread easily in populations of the honey bee parasite *Varroa destructor*

**DOI:** 10.1101/2021.04.30.442149

**Authors:** Nicky R. Faber, Adriaan B. Meiborg, Gus R. McFarlane, Gregor Gorjanc, Brock A. Harpur

## Abstract

Varroa mites (*Varroa destructor*) are the most significant threat to beekeeping worldwide. They are directly or indirectly responsible for millions of colony losses each year. Beekeepers are somewhat able to control Varroa populations through the use of physical and chemical treatments. However, these methods range in effectiveness, can harm honey bees, can be physically demanding on the beekeeper, and do not always provide complete protection from Varroa. More importantly, in some populations Varroa mites have developed resistance to available acaricides. Overcoming the Varroa mite problem will require novel and targeted treatment options. Here, we explore the potential of gene drive technology to control Varroa. We show that spreading a neutral gene drive in Varroa is possible but requires specific colony-level management practices to overcome the challenges of both inbreeding and haplodiploidy. Furthermore, continued treatment with acaricides is necessary to give a gene drive time to fix in the Varroa population. Unfortunately, a gene drive that impacts female or male fertility does not spread in Varroa. Therefore, we suggest that the most promising way forward is to use a gene drive which carries a toxin precursor or removes acaricide resistance alleles.

## Introduction

When the Varroa mite *(Varroa destructor)* jumped from its original host the Eastern honey bee *(Apis cerana)* to the Western honey bee *(Apis mellifera),* it spread rapidly around the globe and caused catastrophic losses of commercial and feral honey bee colonies (1–4). To this day, Varroa mites remain the most highly-reported cause of colony loss for commercial beekeepers and hobbyists (1, 5–7). There are treatment options available to beekeepers that allow them to control Varroa. Unfortunately, currently available treatments do not provide complete protection from Varroa and they often harm honey bees or are physically demanding for the beekeeper. For example, acaricides are among the most effective treatments available and can kill between 49-82% of the Varroa within a colony (8–10). Despite their effectiveness, some acaricides also affect honey bees; they reduce honey bee fertility (11), foraging, and immune responses against bacterial infections (12). More concerning still, in some populations Varroa mites have developed resistance to acaricides (13–16). Beyond chemical treatments, beekeepers can use physical means of Varroa control such as drone brood removal, which gives Varroa mites limited opportunities to reproduce. However, physical methods can require significant labour and thus may not be feasible on a large scale (17, 18). The unfortunate fact of Varroa mite control is that it relies on blunt chemical treatment methods that can harm bees and may not be effective long-term because of evolved resistance. This echoes similar treatment methods available to other pest species around the globe like malarial-vectoring mosquitoes and crop pests like spider mites (19–22).

Genetic population controls, like those that can be implemented through the use of a gene drive (23), could be a more successful and more sustainable means to control Varroa mites and other invertebrate pests than currently-available chemical and physical methods (24). Gene drives are selfish genetic elements that can be engineered to promote the inheritance of desired alleles at rates much greater than conventional Mendelian inheritance (25). When a gene drive allele is introduced into a population, it spreads through the mating of gene drive carrying individuals with wild-type individuals (24). A CRISPR-based gene drive element encodes the two components of CRISPR (a Cas nuclease and guide RNA) and can contain a gene of interest one wishes to propagate (26, 27), or it can be targeted to a gene one wants to disrupt (28–30). In the germline of gene drive carriers, the Cas nuclease and guide RNA are expressed to generate a doublestranded DNA break on the opposing wild-type chromosome at the gene drive locus. This DNA break is repaired through homology-directed repair, using the gene drive harbouring chromosome as the repair template, and thus the gene drive element is copied to the second chromosome (24). The conversion rates for gene drives in insects can be as high as 100% (26, 31–33). This process occurs again in the offspring generation and will do so in all subsequent generations, resulting in the gene drive spreading through the target population. A gene drive can be designed to reduce the fitness of individual homozygous carriers with the aim to reduce population size or even achieve extirpation (23, 34).

The introduction of CRISPR-Cas9 gene drives as a management tool for Varroa numbers could greatly impact our ability to control them, and technology is progressing to a stage where we could test this strategy. The necessary biochemical and biological research is currently coming together: *in vitro*-rearing techniques for Varroa are being refined (35, 36), there is a high-quality reference genome (37), and there is a growing list of genes essential to mite survival (38). CRISPR-Cas9-mediated mutagenesis has not yet been published for Varroa mites but recent work on spider mites demonstrates that this may soon be possible (39). However, we do not yet know if a gene drive can spread in a Varroa population. Prior to any gene drive system being implemented, it is essential to develop a species-specific genetic and demographic model to predict the effectiveness of a drive spreading successfully (29, 30, 34, 40–44). This is especially important in non-model species where mating biology and sexdetermination systems can limit the spread of gene drives. In the case of Varroa mites, they can both outbreed and inbreed, and the proportion of each breeding strategy varies throughout the season based on brood cell availability (44, 45). Inbreeding, along with haplodiploidy (46) in Varroa reduce the likelihood of a gene drive spreading effectively.

We present a modelling study to investigate the effectiveness of a gene drive given the unique life history of Varroa. We estimate the spreading efficiency of a gene drive in a single honey bee colony and identify management techniques beekeepers may have to implement to successfully spread a gene drive in their colonies. We show that spreading a neutral gene drive in Varroa is challenging because of the high rate of inbreeding and their exponential growth rate that can quickly overwhelm a honey bee colony. Some management strategies, including the use of acaricides, may help spread gene drive alleles. Unfortunately, we could devise no scenario to spread gene drives that impact fitness traits like male or female fertility. Therefore, we suggest that the most promising way forward is to use a gene drive which carries a toxin precursor or removes acaricide resistance alleles.

## Results

### A. Development of a genetic population model of *Varroa destructor*

We first created a realistic, stochastic, population model of *Varroa destructor* that includes genetic inheritance. For an overview and description of the model and life history parameters, see Figure 1 and Methods. Our model has a population trajectory that is similar both in shape and amplitude to previous modeling (47–49) and empirical studies (50) (Figure 2A). The model begins on day 1 of the calendar year, a period of low or no growth for temperate populations. The population steadily declines due to daily mortality. By the summer, the Varroa population grows exponentially. The starting population of Varroa greatly influences the speed with which Varroa reach threshold levels within a colony. With 100, 10, or 1 initial Varroa, it respectively takes one, two, or three years longer for the population to reach the threshold of 10,000 individuals where we stop our model. The level of Varroa infestation at which beekeepers will typically treat colonies is reached a year earlier. With 1 initial Varroa, this single Varroa often dies in the winter and therefore, the population grows in only a small number of replicates. Importantly, we observe more variability in models that begin with fewer Varroa. This variability is caused by the timing of reproduction of few Varroa, where small initial differences will grow bigger with the exponential growth.

**Figure 1.**
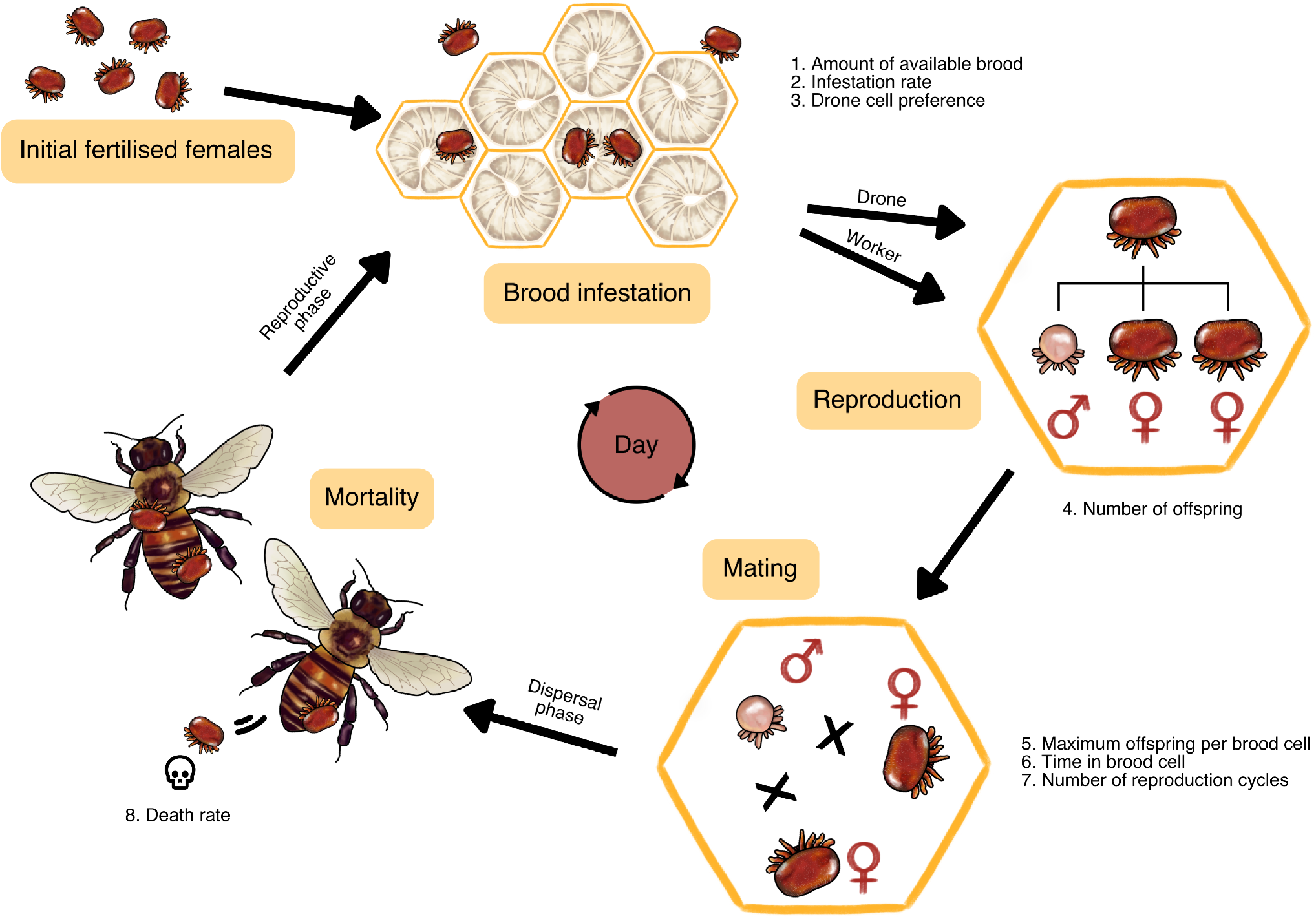
An overview of our Varroa demographic model. For full details, see the Methods section. First, we initialise a certain number of fertilised females. Then, we use a backbone model of an average honey bee colony in a temperate climate where a certain amount of new brood cells become available for Varroa infestation every day. The Varroa infest these cells at a certain rate depending on the number of brood cells and adult bees. Varroa prefer drone cells over worker cells, because those are capped for 2 days longer (14 instead of 12 days), which enables more Varroa offspring to mature. Once in the cell, the fertilised females lay 1 male offspring followed by a varying number of female offspring. Once the females mature, they mate with the male. We assign each female a certain number of reproduction cycles, so one Varroa female can infest brood cells multiple times throughout her life. Then, the fully grown bee emerges from the cell with the Varroa attached to them, which is the start of the Varroa’s dispersal phase. At this stage we model a certain mortality rate which accounts for all ways in which a Varroa could have died during its life cycle.

We were also able to quantify the seasonal fluctuations in inbreeding in our modelled population (Figure 2B). We estimated the mean homozygosity at 1000 bi-allelic loci (with an initial average allele frequency of 0.5) across a single recombining chromosome. We began each model with a mean homozygosity at the beginning of the year of 0.95 in line with previous estimates for Varroa (51). We found that homozygosity remains high throughout most of the beekeeping season but there are pronounced drops in homozygosity during the end of a typical year. This represents a period of time when honey bee colonies are reducing brood production and Varroa populations are typically high. This combination increases the amount of mated Varroa sharing cells, increases the chance of their offspring outbreeding, and thus reducing homozygosity. Overall, our model is qualitatively similar to expectations for a typical Varroa population in a managed honey bee colony living in a temperate climate.

**Figure 2.**
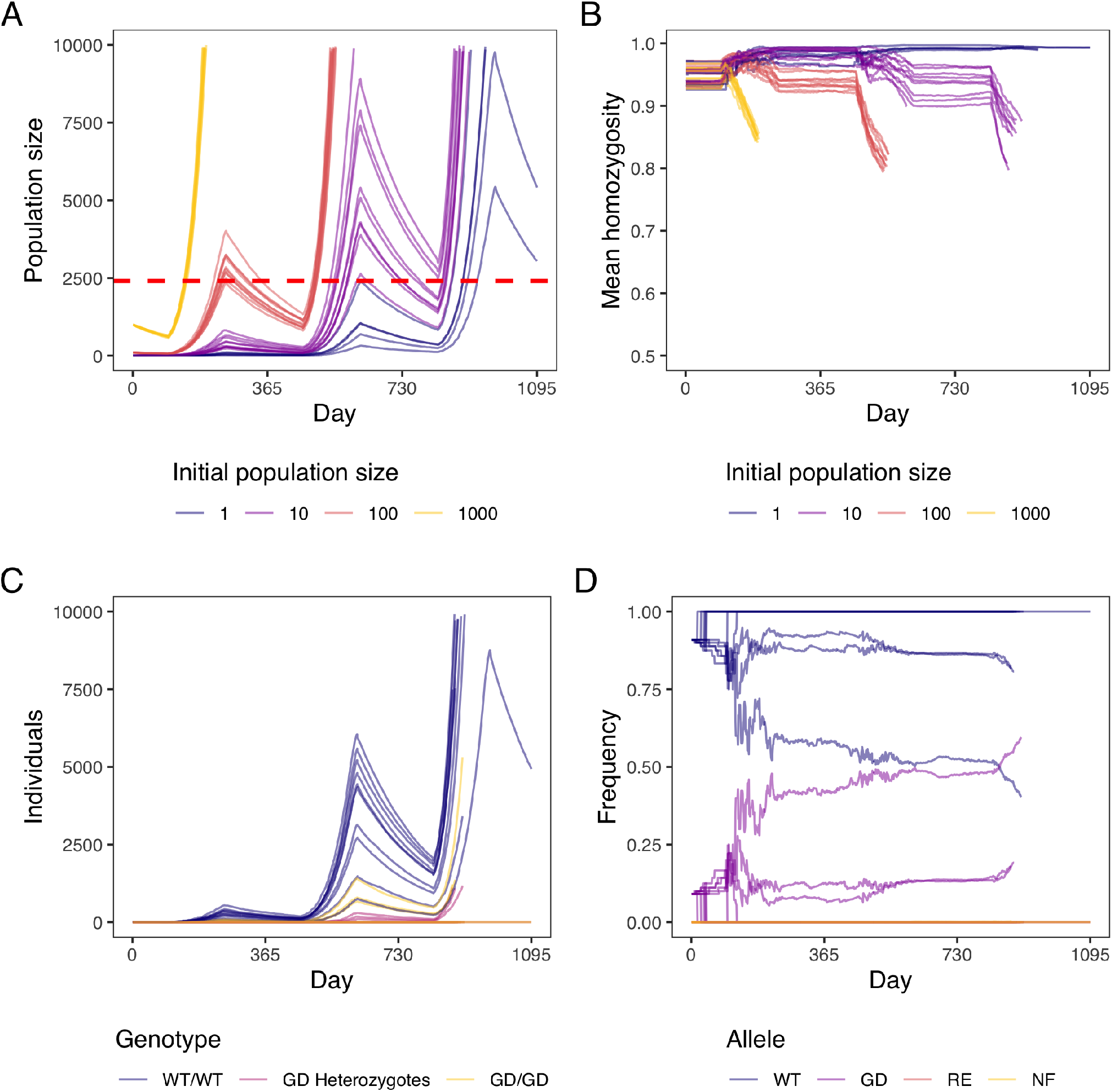
Model of Varroa and gene drive spread. For every set of parameters, we run 10 repetitions and stop the model when the Varroa population size is over 10,000. **A)** Population size over three years with different initial population sizes. The dashed red line indicates a Varroa prevalence of 5% in summer (5 Varroa per 100 adult bees), which is used by beekeepers as a “danger threshold” where treatment is necessary for bee colony health. **B)** Mean homozygosity over three years with different initial population sizes. We model a single chromosome with 1000 bi-allelic loci, each with initial average frequency of 0.5. We initiate individuals at 95% homozygosity because Varroa have very high inbreeding coefficients of 0.9. **C)** Numbers of individuals with three genotypes over three years: WT = wild-type and GD = gene drive. The initial population size was 10 wild-type Varroa with 1 added homozygous gene drive Varroa. **D)** Frequencies of gene drive alleles over three years: WT = wild-type, GD = gene drive, RE = resistant, and NF = non-functional. The initial population size was 10 wild-type Varroa with 1 added homozygous gene drive Varroa, giving an initial gene drive frequency of 0.09.

### B. Inbreeding hinders gene drive spread and a fitness-affecting gene drive cannot spread

We model the release of 1 homozygous gene drive carrying Varroa into a population of 10 wild-type Varroa (gene drive frequency of 0.09), which is relatively high for a non-threshold dependent gene drive (42, 52). We then track the genotypes and allele frequencies of individual Varroa in a single honey bee colony (Figure 2C, D). As can be seen in both plots, the wild-type allele and wild-type genotypes remain the most prevalent even if we allow the model to continue to a population size of 10,000 Varroa mites, greatly exceeding population sizes observed in typical colonies (53). Our model strongly suggests that typical gene drive release frequencies may not be sufficient to spread a gene drive in Varroa. This is likely a result of inbreeding, given that gene drive homozygotes are more prevalent than gene drive heterozygotes over the course of the simulation (Figure 2C). As well, gene drive alleles only meaningfully increase in the last days of the model when Varroa numbers are high and cell sharing increases. The dynamics described above are consistent even when increasing the initial population size and released gene drive individuals (Figure S1). We found that our model is not sensitive to parameters influencing the spread of gene drive alleles (Figure S2). In the context of population control, the goal of a gene drive is to reduce population sizes by spreading alleles that reduce fitness. We could not conceive a model that successfully spread a male- or female-specific fitness-reducing drive (Figure S3).

### C. With high introduction frequencies, a gene drive approaches fixation

When Varroa numbers are still low at the start of the year, it is possible to introduce a larger amount of gene drive Varroa to immediately obtain a high gene drive allele frequency. More importantly, this higher gene drive allele frequency could ensure that whenever outbreeding occurs, a gene drive Varroa is likely involved. Therefore, we modelled a population of 10 wild-type Varroa with either 1, 10, or 50 added homozygous gene drive Varroa. These amounts respectively give initial gene drive frequencies of 0.09, 0.50, and 0.83. We find that the gene drive allele increases most rapidly at an initial release frequency of 0.5, because an outbreeding event is most likely between a gene drive Varroa and a wild-type Varroa, rather than between two wild-types or between two gene drives (see Figure 3 and Figure S4). Naturally, a high initial gene drive frequency results in the highest gene drive allele frequency in the end. Therefore, a high initial release frequency might be beneficial to spread a gene drive through a Varroa population. Unfortunately, we also see that with an initial amount of 50 gene drive Varroa, the population reaches 10,000 individuals a year sooner than with 1 or 10 added Varroa (see Figure 3).

**Figure 3.**
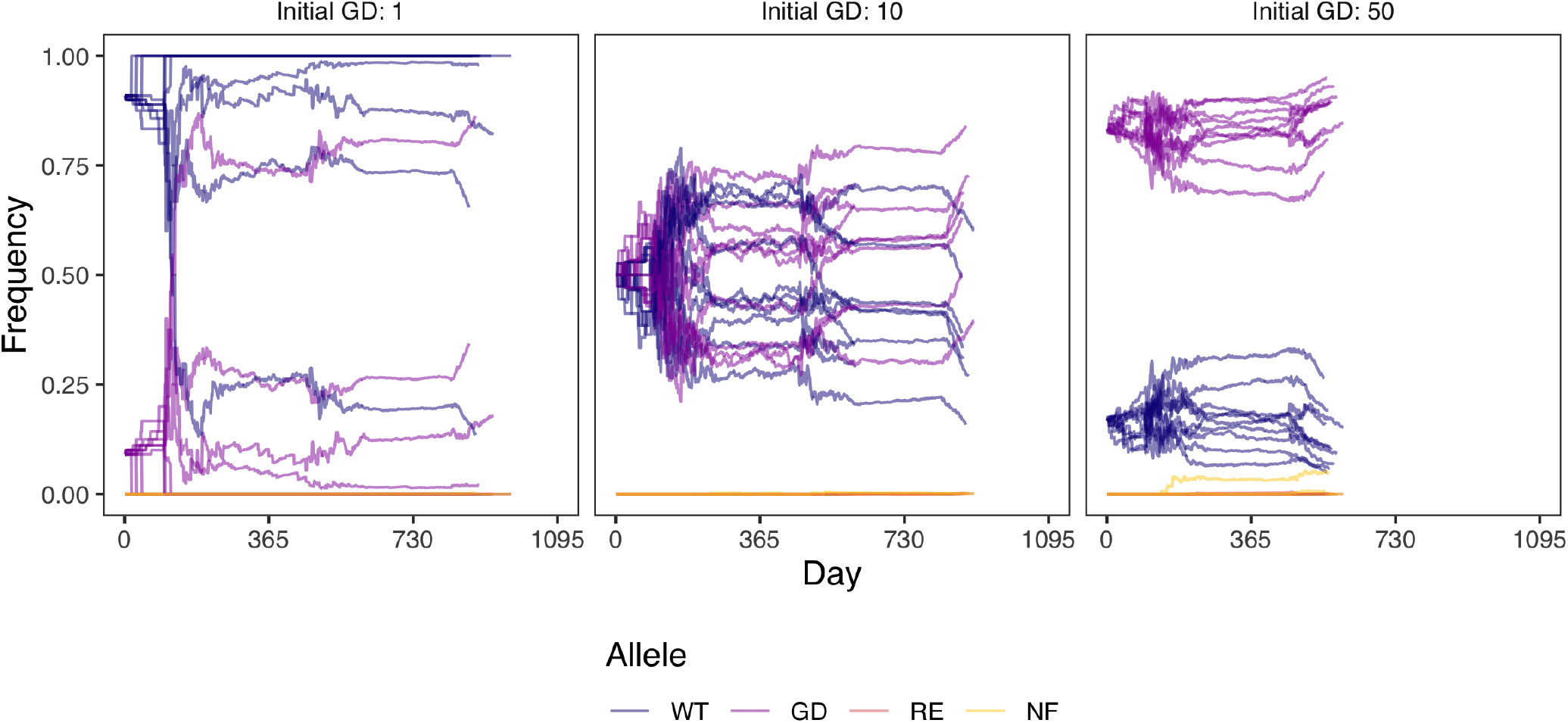
Allele frequencies over three years with different gene drive introductions. The initial population size is 10 wild-type Varroa with 1, 10 or 50 added homozygous gene drive Varroa, giving respective initial gene drive frequencies of 0.09, 0.50, and 0.83. WT = wild-type, GD = gene drive, RE = resistant, and NF = non-functional. For every set of parameters, we run 10 repetitions and stop the model when the Varroa population size is over 10,000.

### D. Brood breaks increase outbreeding, but do not meaningfully increase the spread of a gene drive

Above, we demonstrate that outbreeding can be impacted by the initial release frequency of gene drive Varroa. Ultimately, the amount of cell sharing, and thus outbreeding, depends on three factors: the amount of Varroa, the amount of available brood, and the amount of adult honey bees (54). Therefore, decreasing the number of available honey bee brood cells can increase outbreeding frequency. Cell availability typically decreases naturally at the end of a beekeeping season when honey bees reduce egg laying. Beekeepers can also artificially change cell availability by preventing or restricting queens from laying eggs, a period called a ‘brood break’ (17).

We tested two brood break strategies for their effectiveness at increasing outbreeding and the fixation rate of gene drive alleles. For the first strategy we entirely stopped brood production, forcing Varroa to stay in the dispersal phase (leftmost column in Figure 4). After this brood break, Varroa would more likely infest newly available brood with multiple Varroa per cell. For the second strategy, we provided a steady but lowered amount of brood throughout the brood break (middle three columns in Figure 4). We also modelled no brood break intervention as a control (right-most column in Figure 4). For each of these strategies, we modelled three different brood break starting days: 110 (early season, when brood production is just starting), 160 (middle season, when brood production is at its maximum), and 210 (late season, just before brood production stops). Both strategies increased the amount of cell sharing (see Figure S6). However, only the strategy where a beekeeper adds in a specific proportion of brood during the break increased the the frequency of heterozygous gene drive Varroa in a colony relative to the control without brood break (see Figure 4). A brood break with a beekeeper allowing between 0.01 - 0.1 of available cells to be used for brood was the most effective. In practice, this equates to approximately one full frame in a ten-frame Langstoth colony. These results suggest that with some finetuning, outbreeding can be increased by the beekeeper and therefore increasing the likelihood of fixing a gene drive.

**Figure 4.**
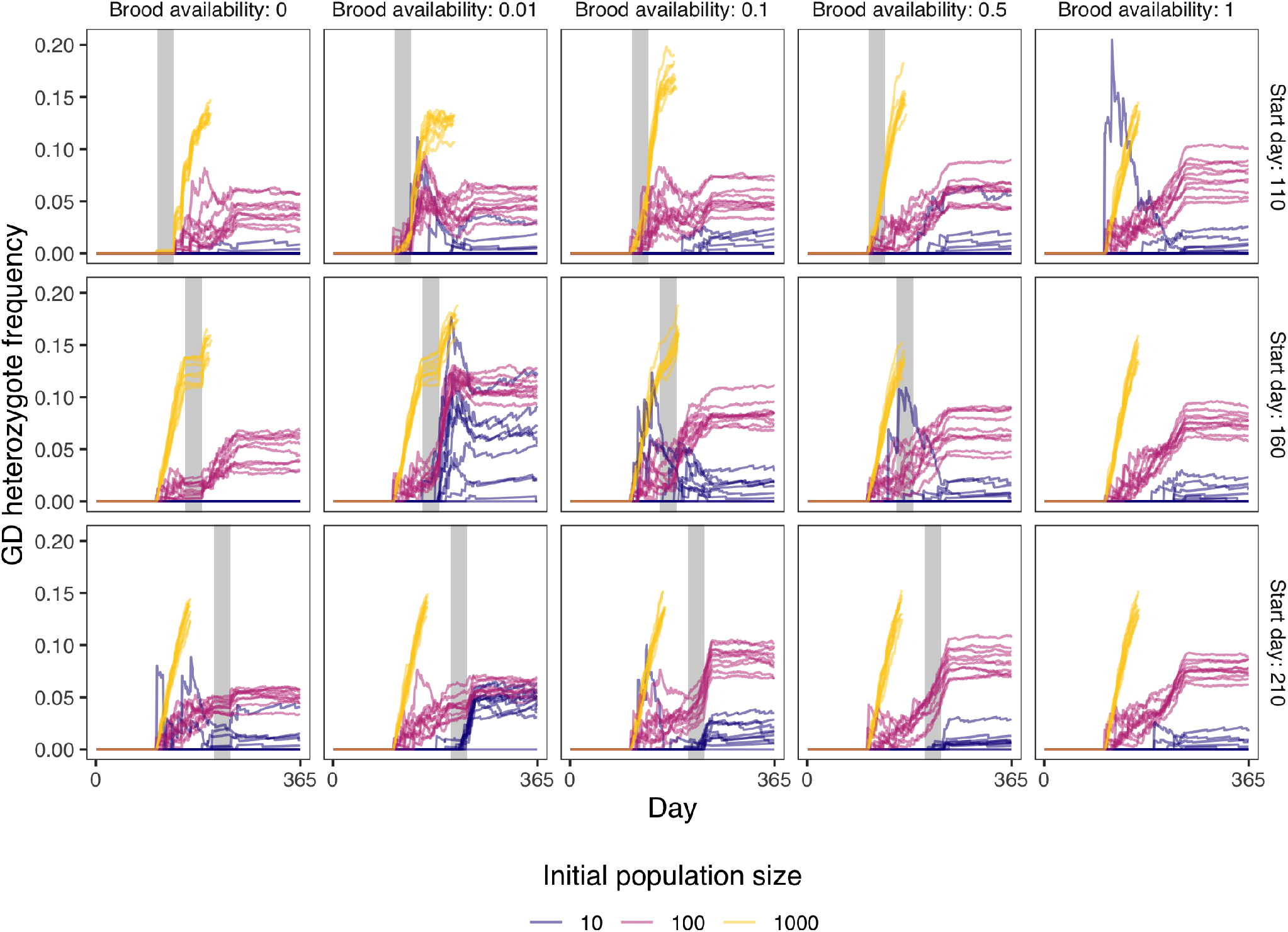
Gene drive (GD) heterozygote frequency over time for different initial population sizes, given different amounts of brood cell availability (as a fraction of the normal amount) and different brood break starting days. The grey bars indicate the brood break. The initial population sizes were 10, 100, or 1000 wild-type Varroa with the same number of gene drive Varroa on top of that, giving an initial gene drive frequency of 0.5. For every set of parameters, we run 10 repetitions and stop the model when the Varroa population size is over 10,000.

Gene drive allele frequency should increase after heterozygotes produce offspring, as gene drive homing will occur in these individuals. Thus, during a brood break, we first expect an increase in heterozygotes as outbreeding occurs, followed by an increase in gene drive allele frequency as these heterozygotes reproduce. However, we show in Figure S7 that there is only a modest increase in gene drive allele frequency after the brood break compared to no brood break. This is likely because of the low frequency of heterozygotes, which is lower than 0.2 as can be seen in Figure 4. In this model, we added the same amount of gene drive Varroa as there are wild-type Varroa, so the allele frequencies are both 0.5. As we showed in Figure 3, this ratio leads to the most rapid increase in gene drive allele frequency. Indeed, in Figure S8 where we model a larger gene drive introduction frequency, the frequency of gene drive heterozygotes is even lower. Despite the high introduction frequency and brood breaks, the gene drive is still not able to fix in the population (see Figure S9). These results show that brood breaks are unlikely to have a large effect on the spread of a gene drive.

### E. Acaricide treatment may facilitate gene drive fixation

None of the scenarios we ran were able to fix a gene drive before Varroa reached threshold levels within a honey bee colony. To that end, we incorporated an acaricide treatment into the model that would be activated anytime a colony reached threshold Varroa levels (Figure 5). We found that effective acaricide treatments provide additional time for a gene drive to reach fixation. However, acaricide treatments significantly increase the variability between the model repetitions, which does not disappear when starting the model with a higher number of initial Varroa (Figure S10). This means that the observed variability is due to the fact that, by chance, we could be removing more gene drive Varroa than wild-types. Therefore, gene drive fixation is not reached very fast and not in all populations.

**Figure 5.**
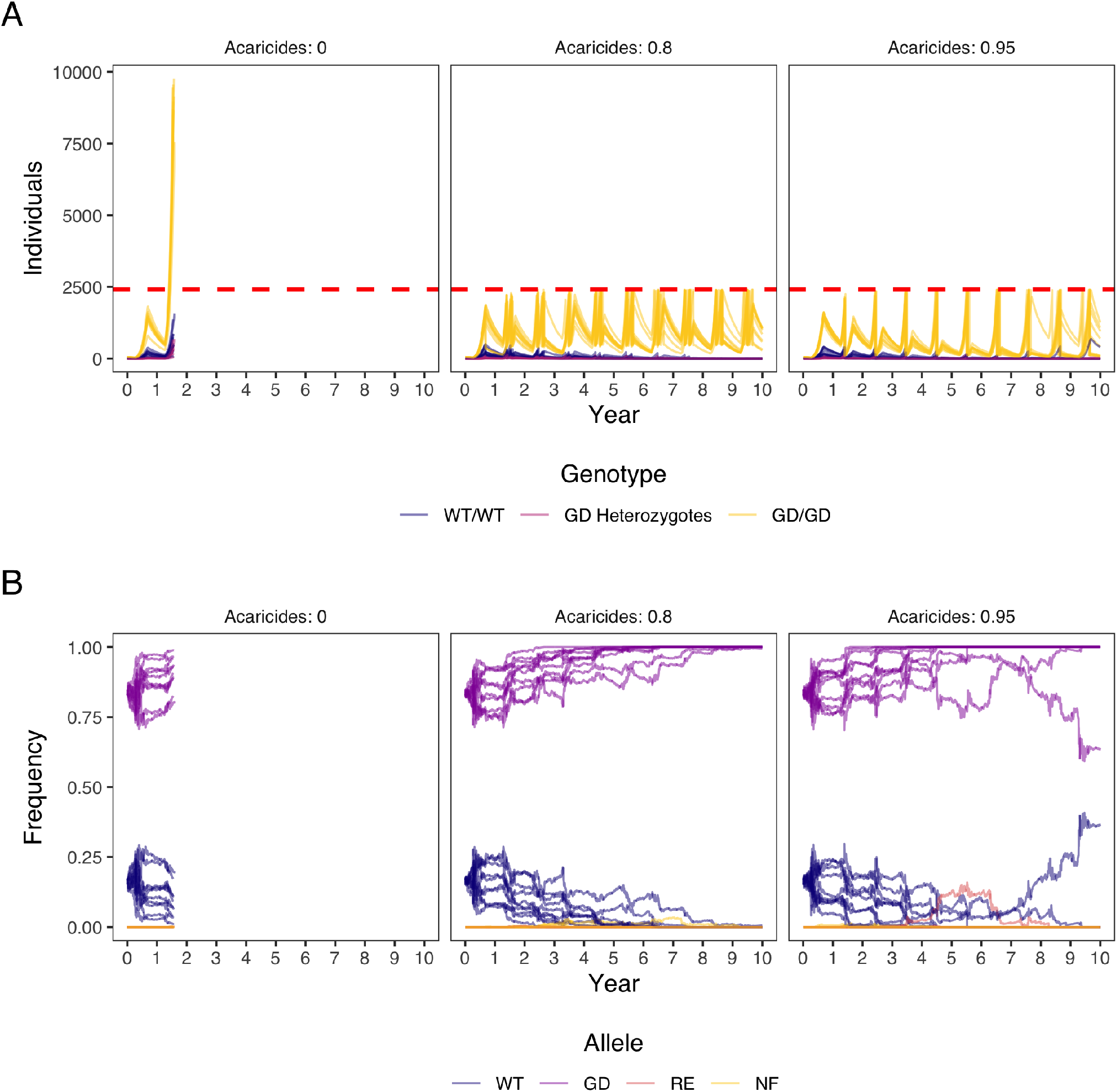
The spread of a gene drive while the Varroa population is suppressed with acaricides whenever the Varroa prevalence surpasses the danger threshold of 5% in summer (5 Varroa per 100 adult bees). The initial population size was 10 wild-type Varroa with 50 homozygous gene drive Varroa, giving an initial gene drive frequency of 0.83. For every set of parameters, we run 10 repetitions and stop the model when the Varroa population size is over 10,000. **A)** Frequencies of gene drive genotypes over time, given different intensities of acaricide treatment when the population surpasses the danger threshold. WT = wild-type, GD = gene drive. **B)** Frequencies of gene drive alleles over time, given different intensities of acaricide treatment when the population surpasses the danger threshold. WT = wild-type, GD = gene drive, RE = resistant, and NF = non-functional.

The best acaricide strategy for gene drive fixation was with 80% acaricide effectivity. With this effectivity Var- roa populations reach the treatment threshold multiple times within a single year and multiple acaricide treatments are necessary. These repeated relatively ineffective treatments are less prone to variability but probably not desirable in practice. We show that introducing more gene drive carriers after acaricide treatment facilitates faster gene drive fixation and less variability (see Figure S11). At this point gene drive fixation is probably due to population replacement rather than gene drive spread.

## Discussion

The greatest threat to managed honey bee colonies, globally, is the Varroa mite (1, 5–7). With the ever-advancing toolkit available to study functional genomics in Varroa (37, 55, 56), we suggest that the prospect of genetic control is not far from a reality. We set out to test the feasibility of such a system, in the form of a gene drive, in a modelling study of a population of Varroa within a single honey bee colony. We demonstrate that a neutral gene drive could spread in a Varroa population in a honey bee colony and open the door to future analysis in exploring how to spread gene drives in non-model species with particularly challenging biology.

A gene drive could work in Varroa but it is slow and requires management inputs. Our stochastic model tracked the growth of Varroa mite populations each day over several years in a typical temperate honey bee colony. Varroa living in colonies in non-temperate climates will likely need additional modelling given the very different demography that honey bees have in these areas (57). We focused on temperate colonies, specifically, because they represent most managed colonies in the United States (5) and because temperate climates provide an opportunity for increased outbreeding in Varroa. Varroa populations tend to be highest in the fall (47, 58, 59). During this time, honey bee colonies decrease brood production to prepare for the winter. As we observe and others have empirically demonstrated, Varroa mites increase outbreeding rates in the fall because of reduced brood cell availability (51). Outbreeding is critical to the establishment of a Varroa gene drive and indeed to any gene drive (45).

We could not conceive a model that would successfully spread a lethal gene drive in Varroa. The most promising way forward may be to design neutral drives with environmentally-induced fitness effects (such as the spreading a toxin precursor), drives which remove acaricide resistance alleles, or drives that target genes involved in Varroa-viral interactions. Each of these requires a deeper understanding of Varroa functional genomics but may be fruitfu for future investigations. Spreading drives that confer Varroa with genetic resistance against viruses is a particularly interesting prospect. The threat that Varroa mites pose to honey bees is exacerbated by the viruses they introduce into their hosts (60–62).

There are several challenges to establishing a gene drive in Varroa that need to be overcome. Natural outbreeding alone was not enough to reliably increase the frequency of gene drive. We attempted to overcome this challenge by incorporating beekeeper management in the form of brood breaks and acaricide treatments. Both influenced the rate of outbreeding and the likelihood of gene drive fixation. Importantly, both of these management practices are used by beekeepers and their incorporation into future gene drive efforts would not be an additional burden. The need for beekeeper management also suggests that a drive has a limited ability to spread beyond the apiary. All gene drive models we attempted faced the additional challenge of concomitantly minimizing population growth. When Varroa populations exceed economic thresholds, honey bee colonies produce less honey and have a higher probability of collapsing (63, 64). Here, we took a very generous threshold of 5 Varroa/100 bees across the year and ran simulations until Varroa reached 10,000 mites in a single colony — a level that would almost never be observed in a managed colony. Furthermore, because Varroa populations grow exponentially, a honey bee colony can only go without Varroa control for a few years at most, depending on the initial infestation level. Controlling Varroa growth with acaricides was an effective means to improve the spread of neutral gene drives by providing more time for the gene drives to fix before the honey bee colony reached 10,000 Varroa. However, this method in itself is troubling because it does not remove the risk of Varroa populations evolving acaricide resistance nor does it remove the risk that some acaricides pose to honey bees. We feel that the addition of management scenarios in our models and others (30) is particularly important for the gene drive literature and a feature that could be overlooked. Incorporating the typical management practices into models and understanding how they impact gene drive dynamics may be an important addition to future work.

In summary, our models provide an early look at how gene drives may act in the Varroa system. They are by no means comprehensive. Varroa occupy a huge range and experience different colony and apiary environments across it. Location- or management-specific models may reveal that gene drives spread more or less successfully.The genetic background of a honey bee colony and a colony’s response to increasing Varroa loads were also not modelled. Both could impact the spread of a gene drive. The population dynamics for Varroa in Varroa-tolerant or resistant colonies is likely different and could impact the spread of a gene drive, perhaps acting like acaricide treatments and providing a longer time for gene drives to spread. Any colony-level responses to increased levels of Varroa parasitism could increase or decrease the likelihood of a drive spreading. We also did not explore dynamics outside of a single honey bee colony and did not explore the risks of modified Varroa establishing in non-target colonies. Varroa mites are as highly mobile as honey bees and more modelling is necessary to understand the roles of drifting, foraging, robbing, and management in spreading gene drives outside of target colonies (65–68). We suggest, given the difficulty we found in spreading drives in a single colony, that the above factors may be unlikely to establish drives in non-target colonies. Even if they could establish outside of target colonies, the spread of gene drive Varroa may not be viewed as a major threat, at least in North America. This may not be the case in other parts of its introduced range. In its native range, *Varroa destructor* can be found in low frequency in *Apis cerana* colonies where we have little information about its native ecology.

To our knowledge, genetic modification has not been performed in Varroa mites and *in vitro* rearing methods are, so far, unable to maintain a breeding population of Varroa (55). Mutagenesis in chelicerates has recently been accomplished (39) but transgenesis has yet to be achieved. Gene drives may be many years off for Varroa. With more expertise developing in the fields of transgenesis and mutagenesis in arthropods, it is likely that we will see experiments in the Varroa system and we hope that our work can help develop ideas about genetic control of this invasive pest species. In the short-term, currently-available treatment methods (63) and perhaps newer methods (38, 69) remain the best methods to control Varroa.

## Methods

Within R 4.0.5 (70), we used the package AlphaSimR as a framework for our modelling (71). AlphaSimR is designed to model the genetics of plant and animal breeding schemes, but lends itself well to general population genetics modelling too. We have created an individual-based, stochastic, day-by-day model of *Varroa destructor* (hereafter simply named Varroa), which consist of three aspects: a static honey bee colony as backbone, a stochastic model of Varroa and its life history, and the implementation of a gene drive. Everyday in the model, we track parameters such as the size of the Varroa population, the levels of inbreeding, and the allele frequencies at the gene drive locus, among others.

### A. Honey bee colony simulation

Varroa is a parasite and depends on its host, *Apis mellifera*, for reproduction. Therefore, to realistically model a population of Varroa, we must also model a honey bee colony. We chose to use a static model for the honey bee colony, as we are primarily interested in the Varroa population and not the interaction between parasite and host. We used a honey bee colony model from Calis et al. (1999) (48), who based their model on data from Allen (1965) (72). This model is based on a colony of average size in a Northern European climate and contains the amount of adult honey bees, drone brood, and worker brood over 365 days. At the end of the year, bee and brood numbers are the same as at the start of the year. Therefore, we can model multiple years by replicating this honey bee model several times back to back. We assumed that a honey bee colony would collapse when the Varroa population reaches 10,000 individuals, at which point we stopped the model. We also implemented an option to reduce brood amounts through colony management by the beekeeper to manage inbreeding in the Varroa population (73). For a variable amount of days, we reduce the brood by a variable percentage of its original amount on those days. In our fixed honey bee colony model, we only change the amount of drone and worker brood and leave the adult bee numbers the same.

### B. Varroa life history

Our model consists of a number of steps to accurately represent the complex life history of Varroa mites:

1. **Initialising mated females.** At the start of the model, we initialise a certain number of mated Varroa females. Then, every time when female Varroa offspring is created, we assign each Varroa a certain number of reproduction cycles it will go through in its life. Current estimates of how many reproduction cycles are completed on average range between 2 to 3 (74, 75). Therefore, we assign each female a number between 1 and 4 randomly, which gives an average of 2.5 reproduction cycles.
2. **Brood infestation.** The first step in Varroa reproduction is the infestation of a honey bee brood cell. For the rate of brood entering, we use a model by Boot et al. (1994) (54), who tested several models to predict this rate. On every day of our model, we calculate the number of infestations *(N_i_)* as:

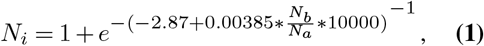

which is dependent on the ratio between available brood *(N_b_)* and the number of adult bees *(N_a_*) (54). The biological reasoning behind this model is that Varroas are phoretic on adults bees and when those bees get close to available brood cells, the Varroa can infest (54). When this ratio is low, the probability that an adult bee with a phoretic Varroa will pass by an available brood cell is low, and vice versa. Once we have determined the number of Varroa that infest, we assign them to the available drone and worker cells. Varroa prefer drone cells over worker cells, because those are capped for 2 days longer (14 instead of 12 days) (47), which enables more Varroa offspring to mature. We model a drone cell preference by giving drone cells an eight times higher probability of infestation (76). Therefore, by chance any drone or worker cell could be infested by more than one Varroa, with the probability of this happening being much higher in drone cells.
3. **Generating offspring.** Varroa mites first produce a single male offspring, followed by a varying number of female offspring (1). More female offspring are able to mature in drone brood than in worker brood because of the longer capping period of those cells (77). Therefore, we use two separate distributions to determine the number of female offspring per Varroa in the two types of brood as described by Infantidis (1984) (59). These distributions include Varroa that produce no offspring as well. The averages of these distributions for female offspring are 1.70 for drone cells and 0.71 for worker cells (59). Excluding the non-productive Varroa, the averages of female offspring are 2.77 for drone cells and 1.33 for worker cells (59).
4. **Mating between offspring.** Varroa offspring mate in the brood cell they are born in (78). Usually only one Varroa infests a cell, which forces offspring to inbreed by full-sibling mating. Occasionally however, especially at the end of the season when Varroa numbers are high, multiple Varroa infest a single cell, which allows for outbreeding (51). Mated females will generate offspring the rest of their lives with the sperm they save in their spermatheca (77). We model random mating between males and females in a brood cell, where females mate with a single male.
5. **Emergence from brood.** In every brood cell, there is a limit to how many Varroa offspring can survive (79). According to data from Martin (1995) (79), the maximum live offspring per cell is 16 in drone cells and 8 in worker cells. Additionally, they show that there is usually one male offspring for every mother mite, so mostly female offspring will not survive in overcrowded brood. This is likely because of competition at the feeding site (79). Therefore, we determine the female offspring survival probability *(P_s_)* per brood cell:

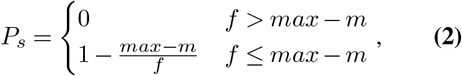

where *(m)* is the number of male offspring, (*f*) the number of female offspring, and *(max)* the maximum number of offspring in that type of brood.
6. **Mortality.** In our model, we expect 0.5% of Varroa to die every day, which is the average between the summer and winter mortality used by Fries et al. (1994) (47). Additionally, we remove Varroa who have gone through their final reproduction cycle, after which they are assumed to die (74).

### C. Gene drive implementation

Although AlphaSimR was designed to model large numbers of loci for breeding and quantitative genetics, the framework is perfect for the single locus of a gene drive too. Each individual is modelled with a single gene drive locus on two chromosomes and inheritance is random.

We have implemented a gene drive which homes in the germline and has four potential alleles: wild-type, gene drive, resistance, and non-functional. Like Prowse et al. (2017) (42), we model a probability of cutting *(P_C_*) of 0.95, a probability of non-homologous end joining (*P_NHEJ_*), which is variable, a probability that non-functional repair occurs (*P_NFR_*) of 0.67, which is the probability of a frame-shift occuring.

## ACKNOWLEDGEMENTS

G.G. acknowledges support from the BBSRC to The Roslin Institute (BBS/E/D/30002275) and The University of Edinburgh’s Data-Driven Innovation Chancellor’s fellowship. B.A.H. was supported by Purdue University and funding from Project Apis M.

## AUTHOR CONTRIBUTIONS

G.G. and B.A.H. conceived the Varroa gene drive project. N.R.F conducted the modelling with assistance from A.B.M. and G.G.. B.A.H. guided the Varroa life history aspects of the project, and G.R.M. and N.R.F. guided the gene drive aspects. N.R.F and B.A.H. wrote the manuscript and all authors reviewed it.

## DATA AVAILABILITY

Our model code and data can be found on the HighlanderLab GitHub: https://github.com/HighlanderLab/nfaber_varroa_gd.

## Supplementary Material

**Figure S1.**
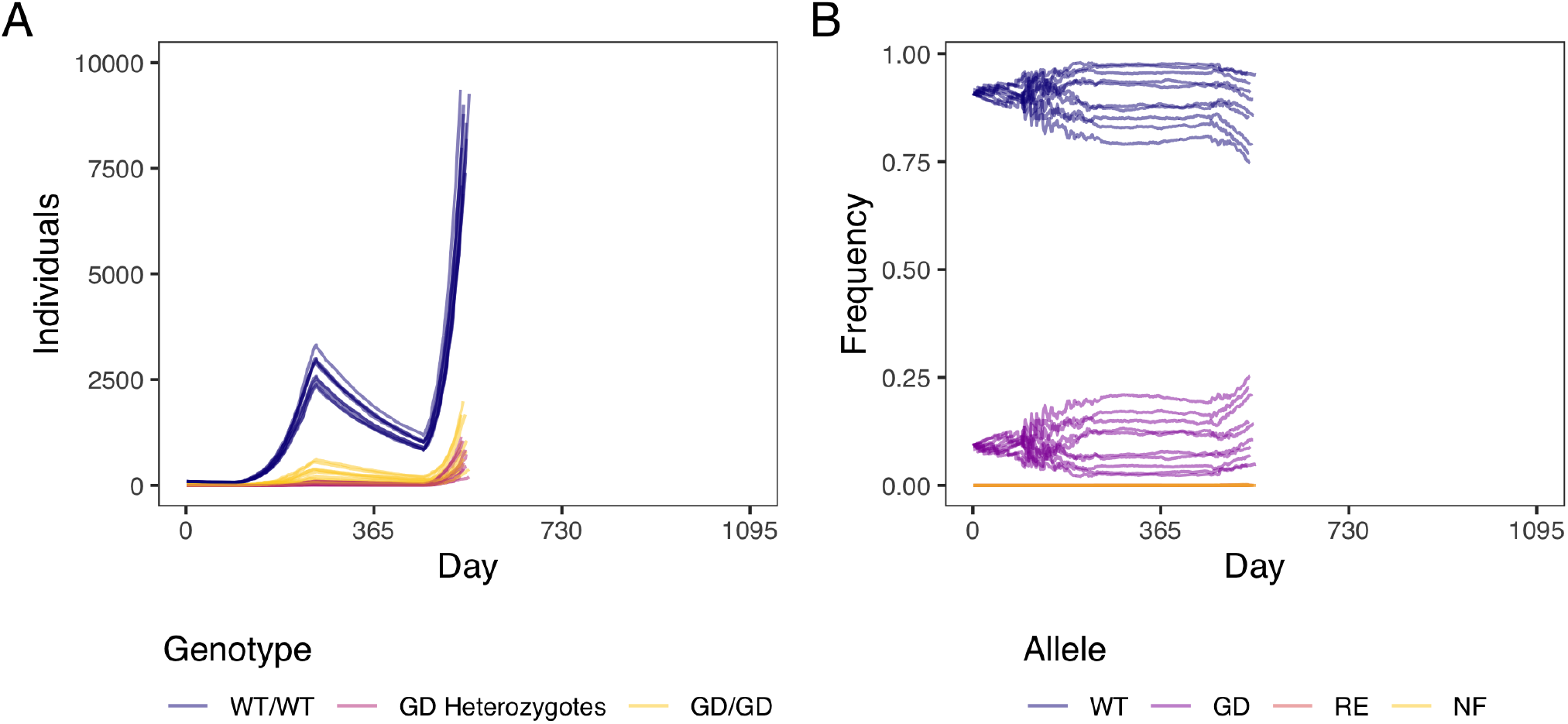
Model of Varroa and gene drive spread as in Figure 2C and D, but with a 10 times larger starting population: 100 wild-type Varroa instead of 10, and 10 gene drive Varroa instead of 1. The initial population size is 100 wild-type Varroa with 10 added homozygous gene drive Varroa, giving an initial gene drive frequency of 0.09. For every set of parameters, we run 10 repetitions and stop the model when the Varroa population size is over 10,000. **A)** Numbers of individuals with different genotypes. WT = wild-type and GD = gene drive. **B)** Frequencies of gene drive alleles. WT = wild-type, GD = gene drive, RE = resistant, and NF = non-functional.

**Figure S2.**
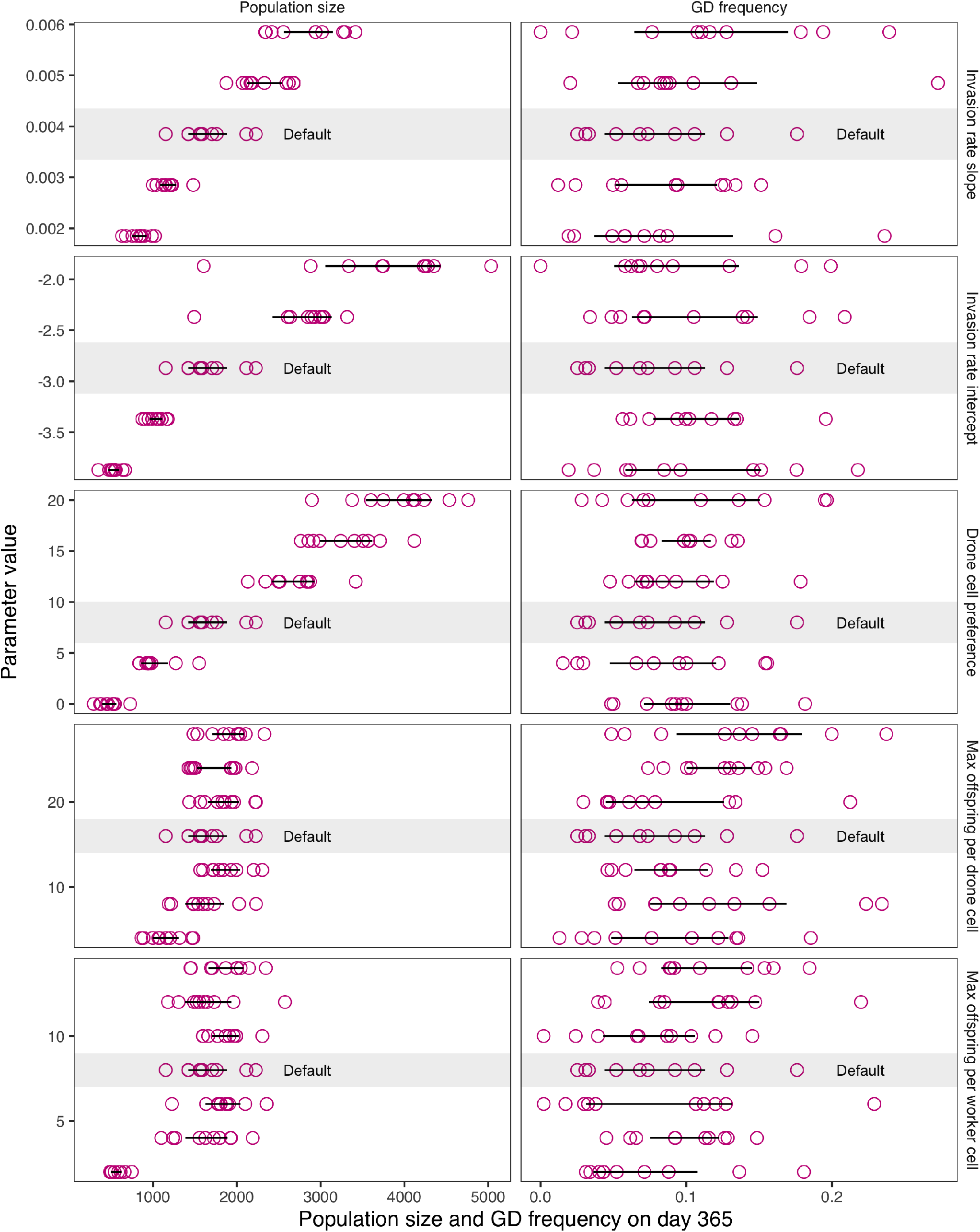
Sensitivity analysis of our Varroa model shown in Figure 2A and D. We run the model for a year with a range of parameters and on day 365, we measure both population size and gene drive (GD) frequency to see which parameter has an influence. The initial population size is 100 wild-type Varroa with 10 added homozygous gene drive Varroa, giving an initial gene drive frequency of 0.09. We vary five parameters independently: invasion rate slope (see Equation 1), invasion rate intercept (see Equation 1), drone cell preference, max offspring per drone cell (see Equation 2), and max offspring per worker cell (see Equation 2). Pink circles indicate each repetition’s outcome, the black lines represent the 95% confidence interval around the mean, and the grey bar and text “Default” indicate the default parameters that are supported by literature and are used in Figure 2 and all other figures. For every set of parameters, we run 10 repetitions and stop the model when the Varroa population size is over 10,000.

**Figure S3.**
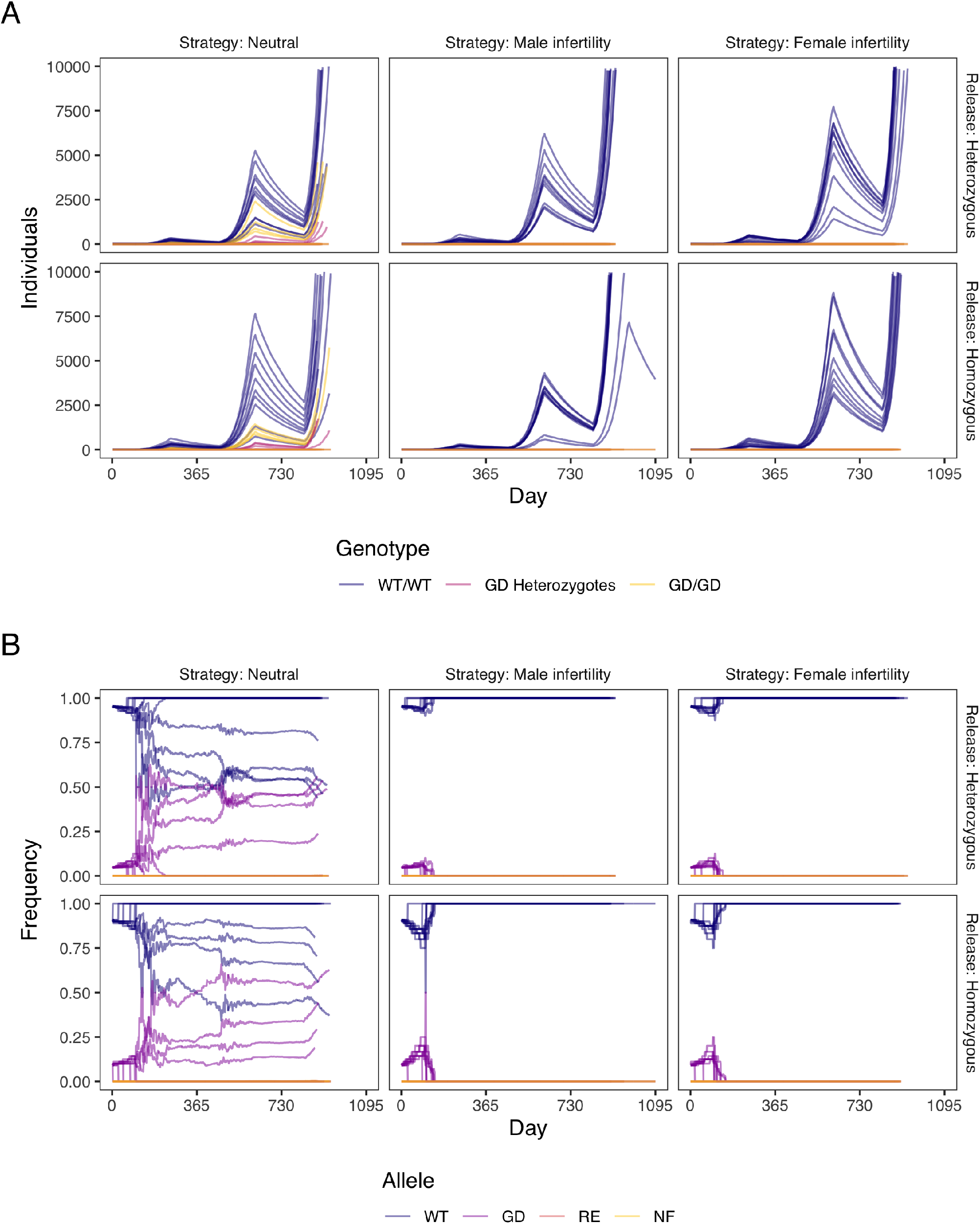
Model of Varroa and gene drive spread as in Figure 2C and D, but besides a neutral gene drive, we also model a gene drive which, when homozygous or hemizygous, causes male or female infertility. Besides the release of homozygous females as in Figure 2C and D, we also model the release of heterozygous gene drive Varroa females so the infertility does not immediately affect females. The initial population size is 10 wild-type Varroa with 1 added gene drive Varroa, giving an initial gene drive frequency of 0.09 for a homozygote release and 0.045 for a heterozygote release. For every set of parameters, we run 10 repetitions and stop the model when the Varroa population size is over 10,000. **A)** Numbers of individuals with different genotypes. WT = wild-type and GD = gene drive. **B)** Frequencies of gene drive alleles. WT = wild-type, GD = gene drive, RE = resistant, and NF = non-functional.

**Figure S4.**
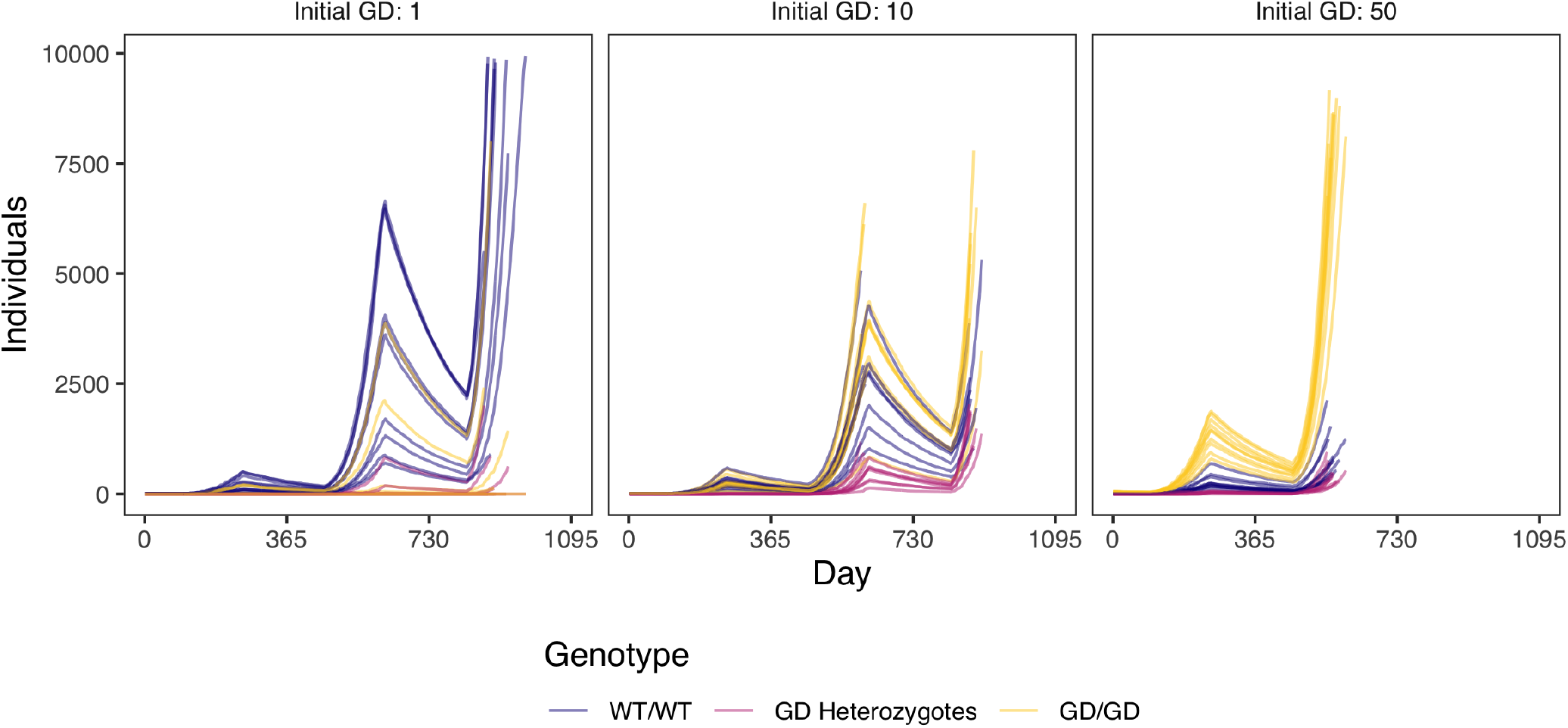
Numbers of individuals with three genotypes, corresponding to the allele frequencies in Figure 3 over three years with different gene drive introduction amounts. The initial population size is 10 wild-type Varroa with 1, 10 or 50 added homozygous gene drive Varroa, giving initial gene drive frequencies of 0.09, 0.50, and 0.83, respectively. WT = wild-type and GD = gene drive. For every set of parameters, we run 10 repetitions and stop the model when the Varroa population size is over 10,000.

**Figure S5.**
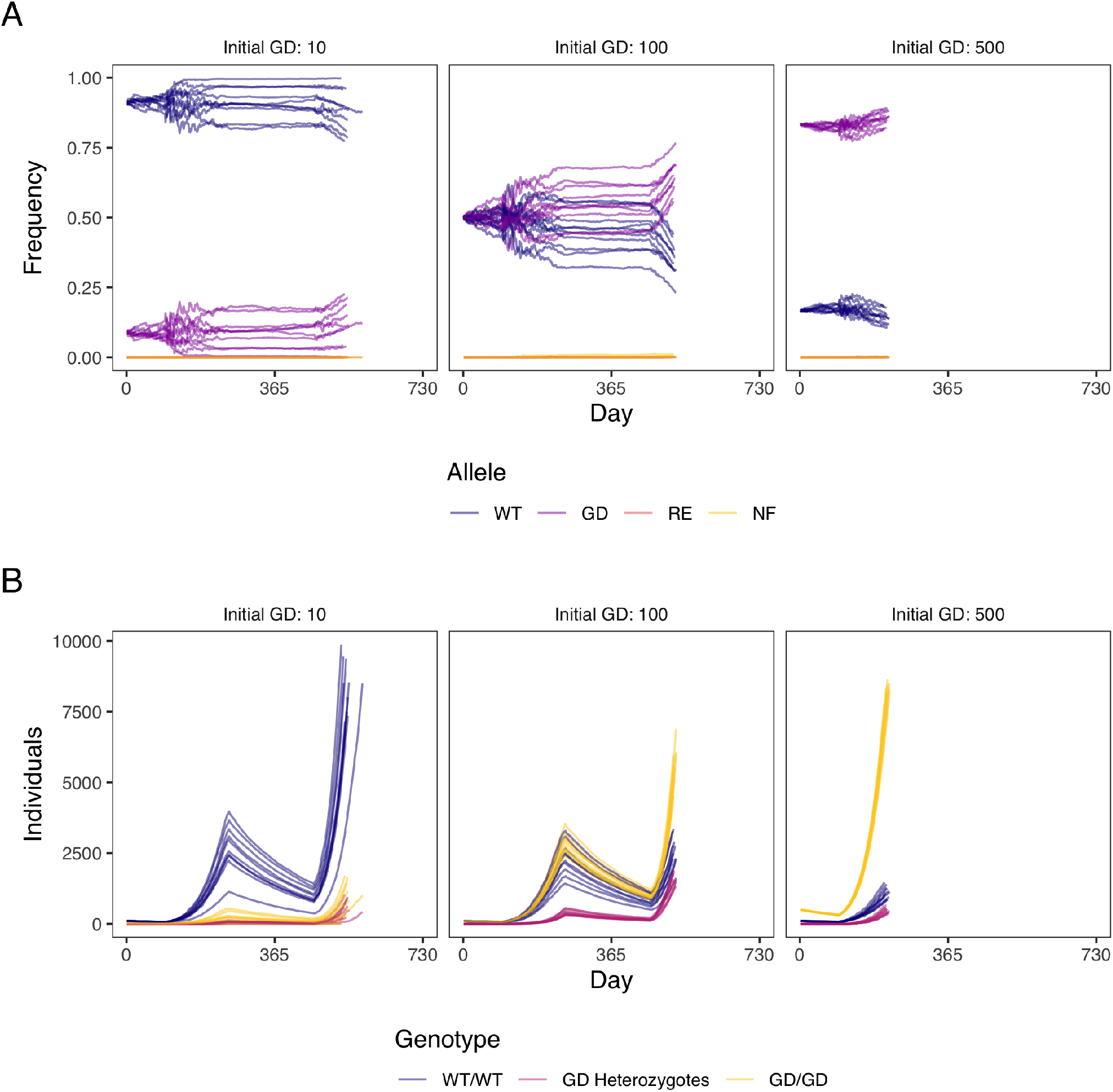
The same as Figure 3 and Figure S4, but with 10 times more initial Varroa. The initial population size is 100 wild-type Varroa with 10, 100 or 500 added homozygous gene drive Varroa, respectively giving initial gene drive frequencies of 0.09, 0.50, and 0.83. For every set of parameters, we run 10 repetitions and stop the model when the Varroa population size is over 10,000. **A)** Allele frequencies over three years with different gene drive introduction amounts. WT = wild-type, GD = gene drive, RE = resistant, and NF = non-functional. **B)** Numbers of individuals with three genotypes. WT = wild-type and GD = gene drive.

**Figure S6.**
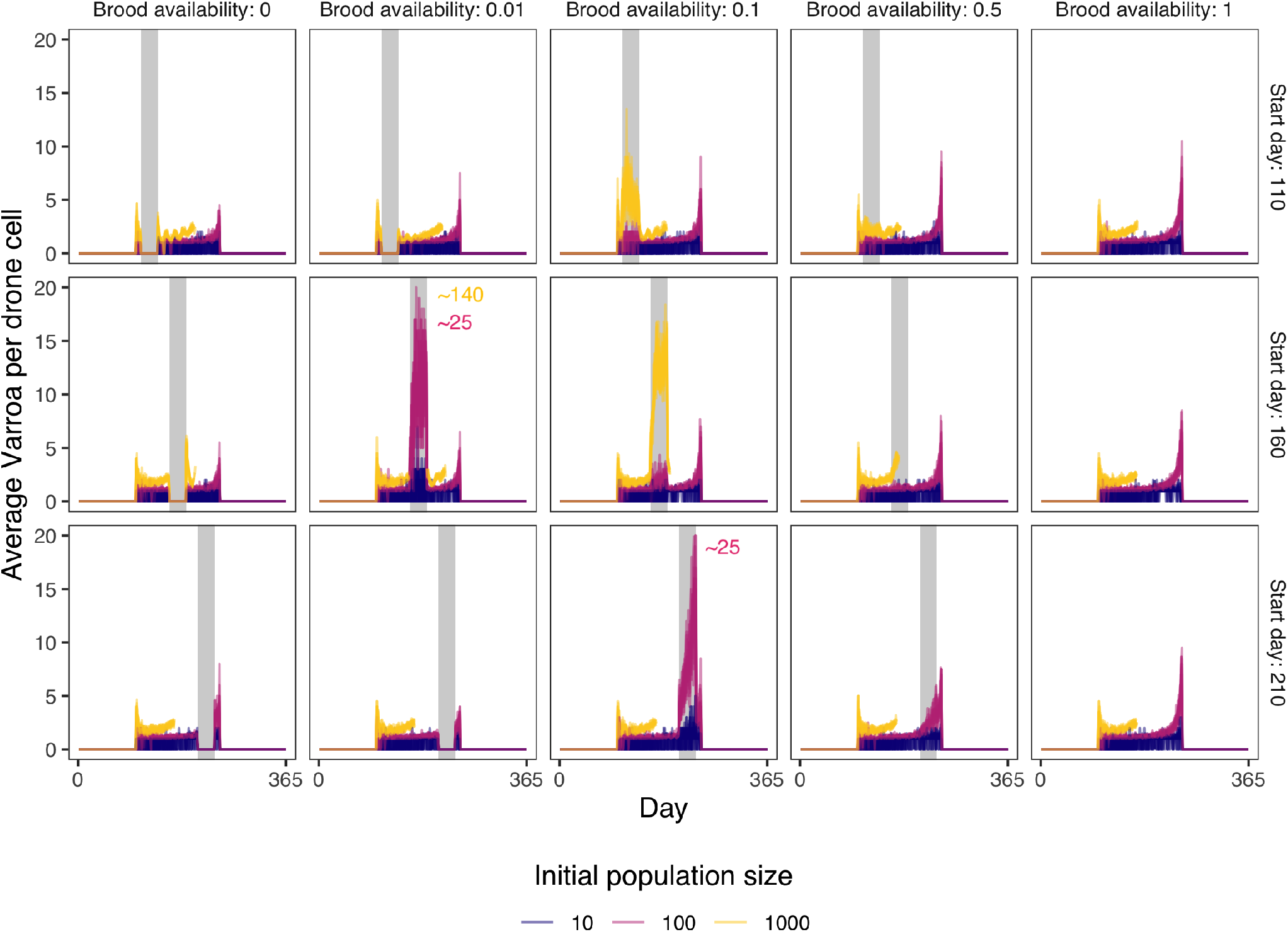
Average Varroa per drone cell over time for different initial population sizes, given different amounts of brood cell availability (as a fraction of the normal amount) and different brood break starting days like in Figure 4. The grey bars indicate the brood break. The “~” in two plots indicates that values were higher than 20 and thus fall off the truncated y-axis to keep the plot interpretable. The number after the “~” roughly indicates the maximum of the truncated values. The initial population sizes were 10, 100, or 1000 wild-type Varroa with the same number of gene drive Varroa on top of that, giving initial gene drive frequencies of 0.5. For every set of parameters, we run 10 repetitions and stop the model when the Varroa population size is over 10,000.

**Figure S7.**
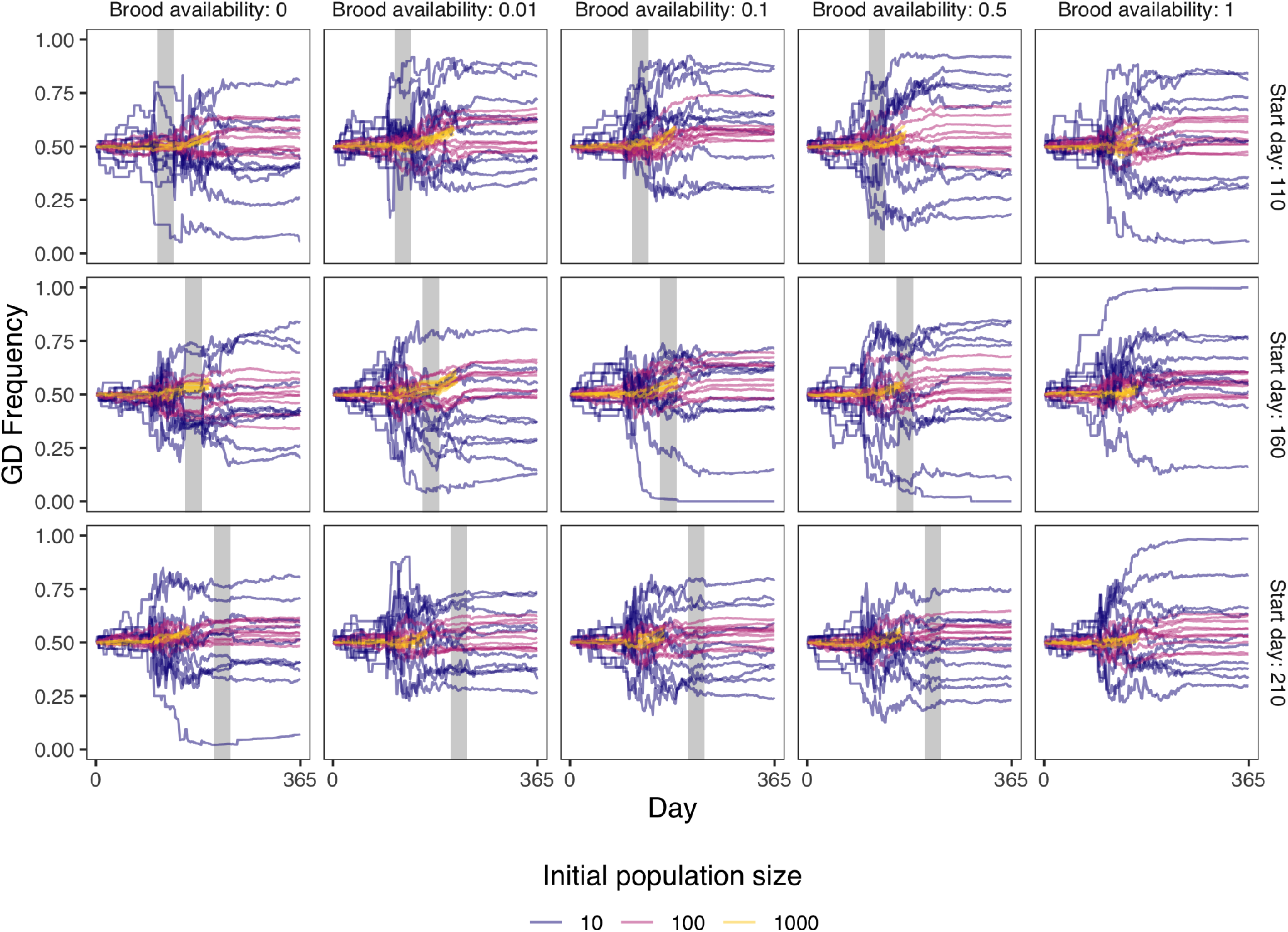
Gene drive (GD) allele frequency overtime for different initial population sizes, given different amounts of brood cell availability (as a fraction of the normal amount) and different brood break starting days like in Figure 4. The grey bars indicate the brood break. The initial population sizes were 10, 100, or 1000 wild-type Varroa with the same number of gene drive Varroa on top of that, giving an initial gene drive frequency of 0.5. For every set of parameters, we run 10 repetitions and stop the model when the Varroa population size is over 10,000.

**Figure S8.**
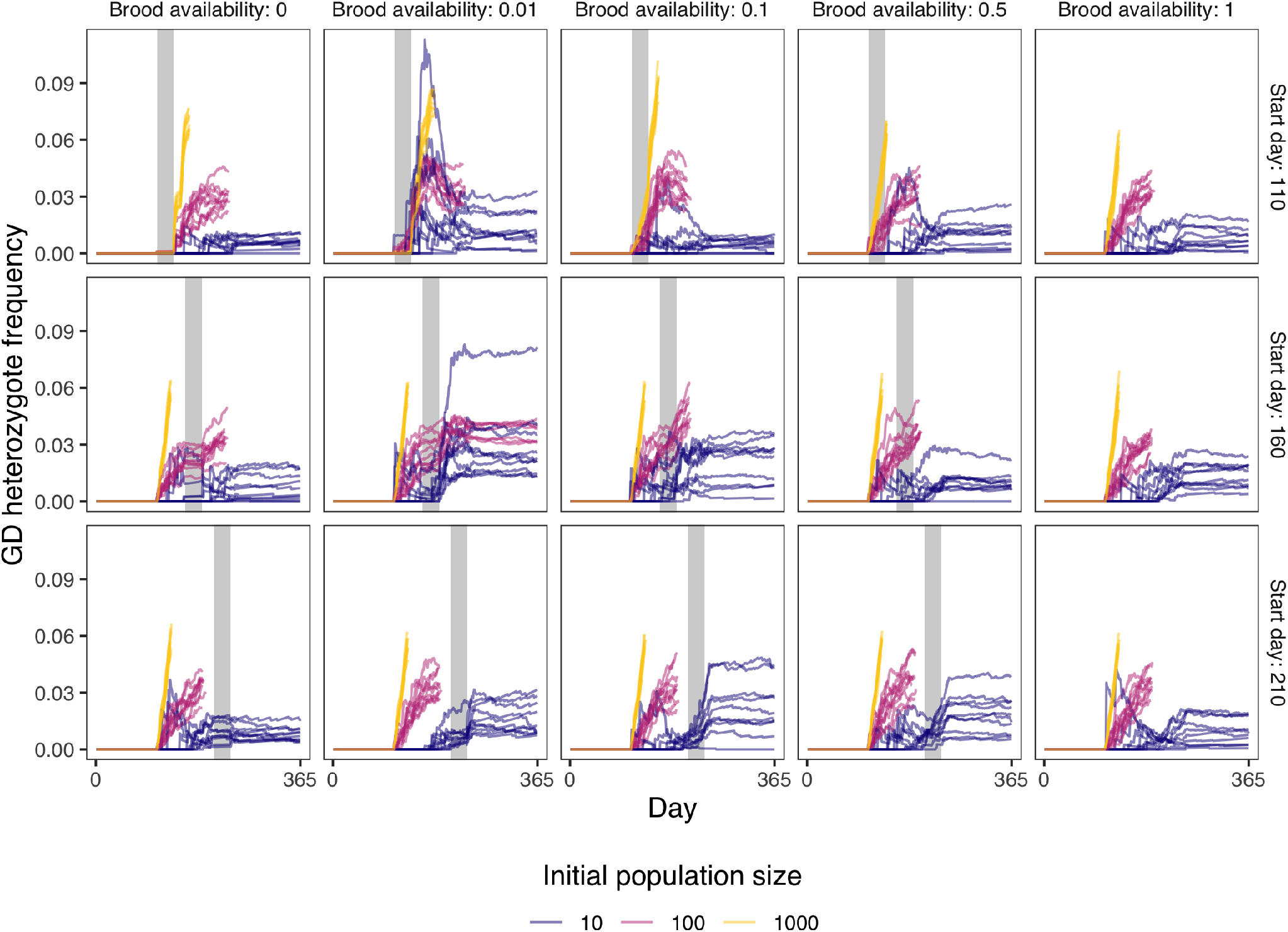
Gene drive (GD) heterozygote frequency over time for different initial population sizes, given different amounts of brood cell availability (as a fraction of the normal amount) and different brood break starting days like in Figure 4, but with more introduced gene drive Varroa. The initial population sizes were 10, 100, and 1000 wild-type Varroa with 100, 1000, and 5000 gene drive Varroa on top of that, respectively, giving initial gene drive frequencies of 0.91, 0.91, and 0.83. The grey bars indicate the brood break. For every set of parameters, we run 10 repetitions and stop the model when the Varroa population size is over 10,000.

**Figure S9.**
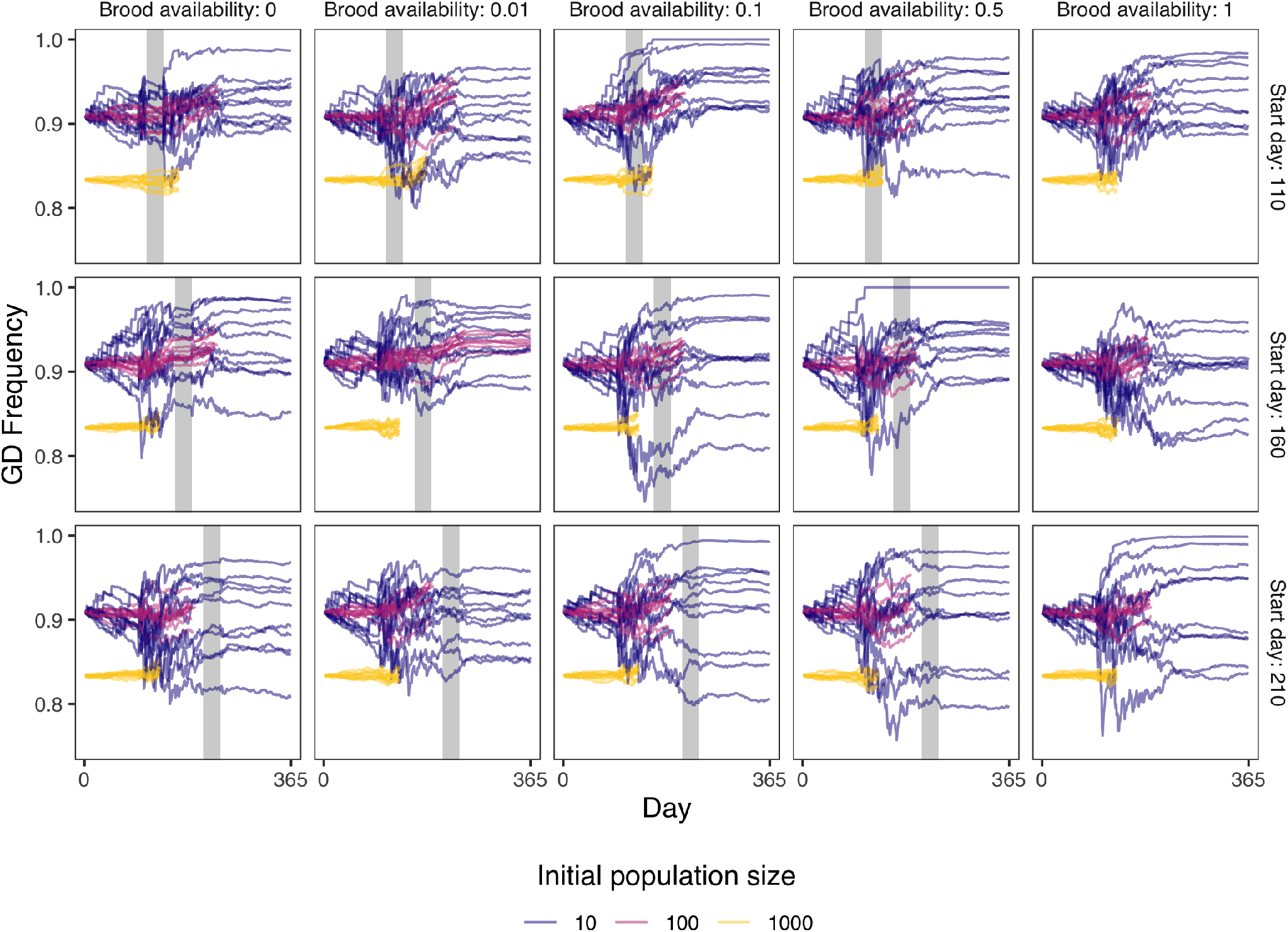
Gene drive (GD) allele frequency overtime for different initial population sizes, given different amounts of brood cell availability (as a fraction of the normal amount) and different brood break starting days like in Figure 4, but with more introduced gene drive Varroa. The initial population sizes were 10, 100, and 1000 wild-type Varroa with 100, 1000, and 5000 gene drive Varroa on top of that, respectively, giving initial gene drive frequencies of 0.91, 0.91, and 0.83. The grey bars indicate the brood break. For every set of parameters, we run 10 repetitions and stop the model when the Varroa population size is over 10,000.

**Figure S10.**
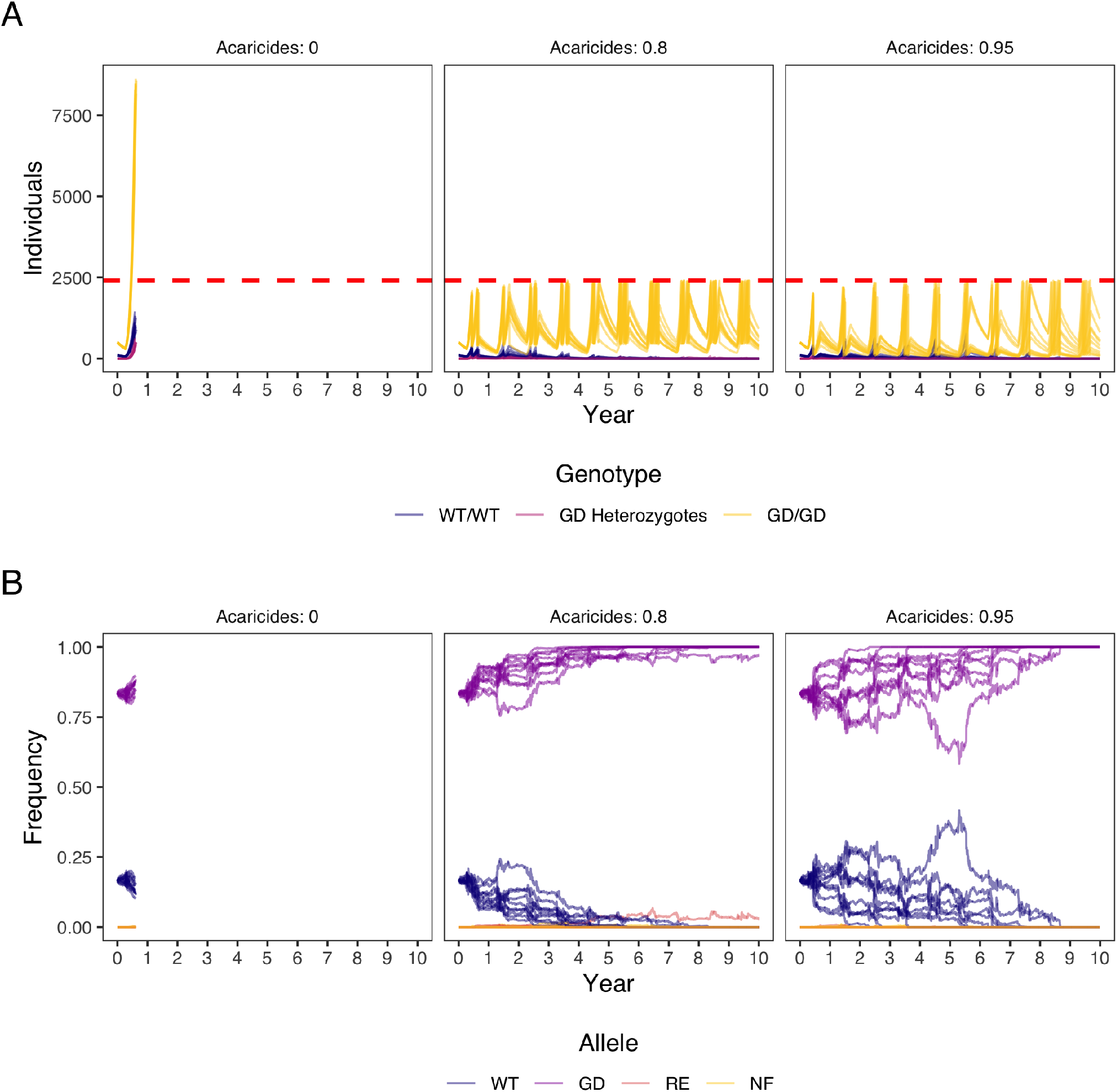
The spread of a gene drive while the Varroa population is suppressed with acaricides whenever the Varroa prevalence surpasses the danger threshold of 5% in summer (5 Varroa per 100 adult bees). The same as Figure 5, but with a 10 times larger starting population. The initial population size was 100 wild-type Varroa with 500 homozygous gene drive Varroa, giving an initial gene drive frequency of 0.83. For every set of parameters, we run 10 repetitions and stop the model when the Varroa population size is over 10,000. **A)** Frequencies of gene drive genotypes over time, given different intensities of acaricide treatment when the population surpasses the danger threshold. WT = wild-type, GD = gene drive. **B)** Frequencies of gene drive alleles over time, given different intensities of acaricide treatment when the population surpasses the danger threshold. WT = wild-type, GD = gene drive, RE = resistant, and NF = non-functional.

**Figure S11.**
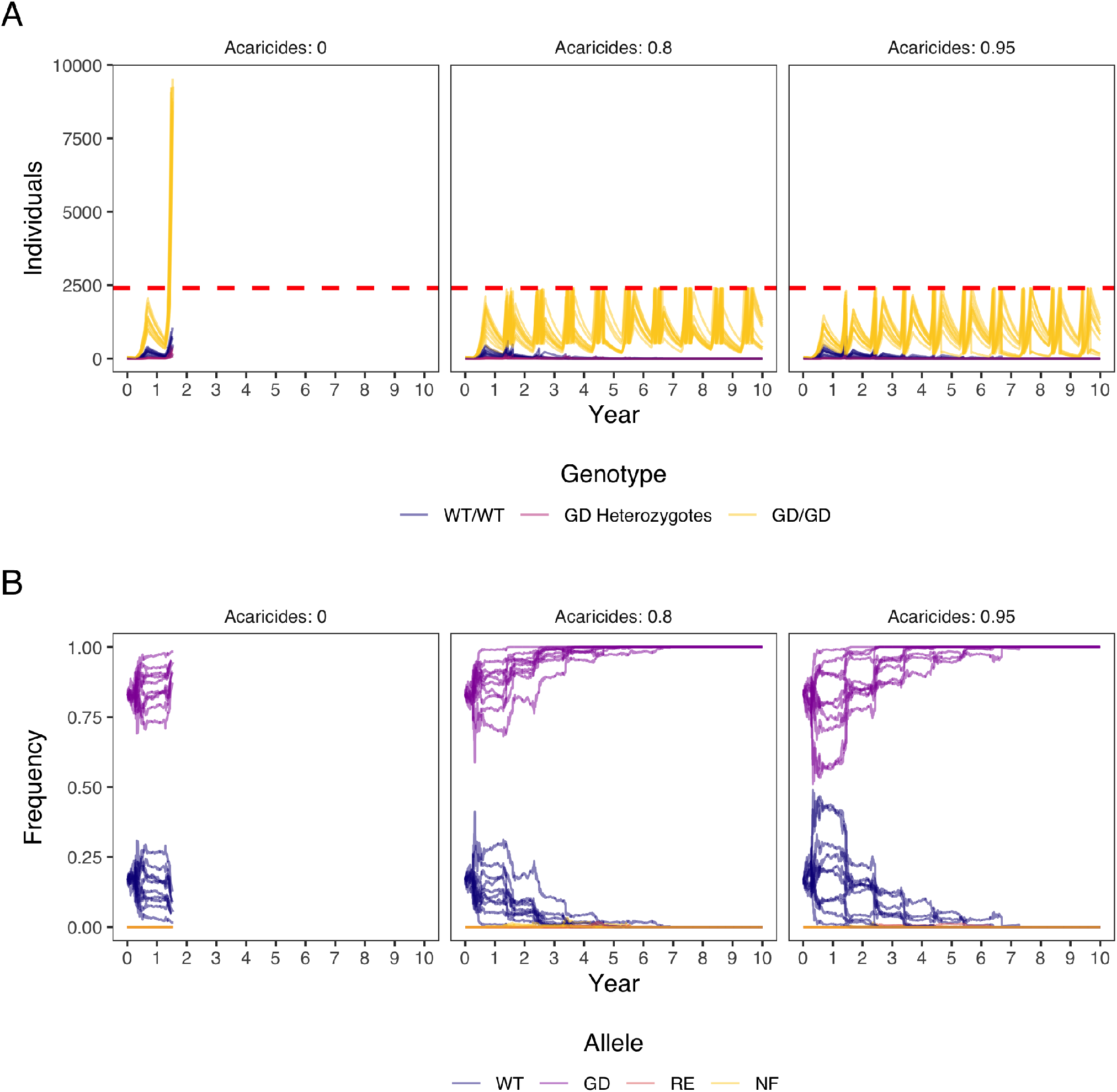
The spread of a gene drive while the Varroa population is suppressed with acaricides whenever the Varroa prevalence surpasses the danger threshold of 5% in summer (5 Varroa per 100 adult bees). The same as Figure 5, but now we do an extra release of 50 gene drive Varroa after every acaricide treatment. The initial population size was 10 wild-type Varroa with 50 homozygous gene drive Varroa, giving an initial gene drive frequency of 0.83. For every set of parameters, we run 10 repetitions and stop the model when the Varroa population size is over 10,000. **A)** Frequencies of gene drive genotypes over time, given different intensities of acaricide treatment when the population surpasses the danger threshold. WT = wild-type, GD = gene drive. **B)** Frequencies of gene drive alleles over time, given different intensities of acaricide treatment when the population surpasses the danger threshold. WT = wild-type, GD = gene drive, RE = resistant, and NF = non-functional.

## References

1. Kirsten S Traynor, Fanny Mondet, Joachim R de Miranda, Maeva Techer, Vienna Kowallik, Melissa AY Oddie, Panuwan Chantawannakul, and Alison McAfee. Varroa destructor: A complex parasite, crippling honey bees worldwide. Trends in Parasitology, 2020.

2. Stephen L Buchmann and Gary Paul Nabhan. The pollination crisis: the plight of the honey bee and the decline of other pollinators imperils future harvests. Sciences, 36:22+, 1996.

3. Adrian M Wenner, W W Bushing, and Others. Varroa mite spread in the united states. Bee Culture, 124(6):341–343, 1996.

4. Bernhard Kraus and Robert E Page. Effect of varroa jacobsoni (mesostigmata: Varroidae) on feral apis mellifera (hymenoptera: Apidae) in california. Environ. Entomol., 24(6):1473–1480, December 1995.

5. Kelly Kulhanek, Nathalie Steinhauer, Karen Rennich, Dewey M Caron, Ramesh R Sagili, Jeff S Pettis, James D Ellis, Michael E Wilson, James T Wilkes, David R Tarpy, Robyn Rose, Kathleen Lee, Juliana Rangel, and Dennis vanEngelsdorp. A national survey of managed honey bee 2015–2016 annual colony losses in the USA. null, 56(4):328–340, August 2017.

6. Dennis vanEngelsdorp and Marina Doris Meixner. A historical review of managed honey bee populations in europe and the united states and the factors that may affect them. J. Invertebr. Pathol., 2010.

7. Ana Molineri, Agostina Giacobino, Adriana Pacini, Natalia Bulacio Cagnolo, Julieta Merke, Emanuel Orellano, Ezequiel Bertozzi, Luis Zago, Andrea Aignasse, Hernán Pietronave, et al. Environment and varroa destructor management as determinant of colony losses in apiaries under temperate and subtropical climate. Journal of Apicultural Research, 57(4): 551–564, 2018.

8. Marco Pietropaoli and Giovanni Formato. Acaricide efficacy and honey bee toxicity of three new formic acid-based products to control varroa destructor. Journal of Apicultural Research, 58(5):824–830, 2019. doi: https://doi.org/10.1080/00218839.2019.1656788.

9. Gerardo Pérez Santiago, Gabriel Otero-Colina, David Mota Sánchez, Martha Elva Ramírez Guzmán, and Rémy Vandame. Comparing effects of three acaricides on varroa jacobsoni (acari: Varroidae) and apis mellifera (hymenoptera: Apidae) using two application techniques. Fla. Entomol., 83(4):468–476, 2000.

10. Morgan A Roth, James M Wilson, Keith R Tignor, and Aaron D Gross. Biology and management of varroa destructor (mesostigmata: Varroidae) in apis mellifera (hymenoptera: Apidae) colonies. J Integr Pest Manag, 11(1), January 2020.

11. Juliana Rangel and Adrian Fisher. Factors affecting the reproductive health of honey bee (apis mellifera) drones—a review. Apidologie, September 2019.

12. Hanan A Gashout, Ernesto Guzman-Novoa, and Paul H Goodwin. Synthetic and natural acaricides impair hygienic and foraging behaviors of honey bees. Apidologie, August 2020.

13. Diana Sammataro, Pia Untalan, Felix Guerrero, and Jennifer Finley. The resistance of varroa mites (acari: Varroidae) to acaricides and the presence of esterase. null, 31(1): 67–74, March 2005.

14. P J Elzen, D Westervelt, and Others. Detection of coumaphos resistance in varroa destructor in florida. Am. Bee. J., 142(4):291–292, 2002.

15. Patti J Elzen, James R Baxter, Marla Spivak, and William T Wilson. Control of varroa jacobsoni oud. resistant to fluvalinate and amitraz using coumaphos. Apidologie, 31(3): 437–441, 2000.

16. Norberto Milani. The resistance of varroa jacobsoni oud. to acaricides. Apidologie, 30(2-3): 229–234, 1999.

17. N W Calderone. Evaluation of drone brood removal for management of varroa destructor (acari: Varroidae) in colonies of apis mellifera (hymenoptera: Apidae) in the northeastern united states. J. Econ. Entomol., 98(3):645–650, June 2005.

18. Nicholas P Aliano and Marion D Ellis. A strategy for using powdered sugar to reduce varroa populations in honey bee colonies. null, 44(2):54–57, January 2005.

19. Rachel Carson. Silent Spring. Houghton Mifflin Harcourt, 1962.

20. Chusak Prasittisuk and James R Busvine. DDT-resistant mosquito strains with crossresistance to pyrethroids. Pestic. Sci., 8(5):527–533, October 1977.

21. Howard Baker. Spider mites, insects and DDT. Yearbook of Agriculture, 1952:562–566, 1952.

22. Timothy J Dennehy, Jeffrey Granett, and Thomas F Leigh. Relevance of Slide-Dip and residual bioassay comparisons to detection of resistance in spider mites. J. Econ. Entomol., 76(6):1225–1230, December 1983.

23. Jackson Champer, Anna Buchman, and Omar S Akbari. Cheating evolution: engineering gene drives to manipulate the fate of wild populations. Nature Reviews Genetics, 17(3):146, 2016. doi: https://www.doi.org/10.1038/nrg.2015.34.

24. Kevin M Esvelt, Andrea L Smidler, Flaminia Catteruccia, and George M Church. Emerging technology: concerning rna-guided gene drives for the alteration of wild populations. Elife, 3:e03401, 2014.

25. Gus R McFarlane, C Bruce A Whitelaw, and Simon G Lillico. Crispr-based gene drives for pest control. Trends in biotechnology, 36(2):130–133, 2018.

26. Valentino M Gantz, Nijole Jasinskiene, Olga Tatarenkova, Aniko Fazekas, Vanessa M Macias, Ethan Bier, and Anthony A James. Highly efficient cas9-mediated gene drive for population modification of the malaria vector mosquito anopheles stephensi. Proc. Natl. Acad. Sci. U. S. A., 112(49):E6736–43, December 2015.

27. Joanna Buchthal, Sam Weiss Evans, Jeantine Lunshof, Sam R Telford, 3rd, and Kevin M Esvelt. Mice against ticks: an experimental community-guided effort to prevent tick-borne disease by altering the shared environment. Philos. Trans. R. Soc. Lond. B Biol. Sci., 374 (1772):20180105, May 2019.

28. Kyros Kyrou, Andrew M Hammond, Roberto Galizi, Nace Kranjc, Austin Burt, Andrea K Beaghton, Tony Nolan, and Andrea Crisanti. A CRISPR-Cas9 gene drive targeting doublesex causes complete population suppression in caged anopheles gambiae mosquitoes. Nat. Biotechnol., 36(11):1062–1066, December 2018.

29. Mohammad KaramiNejadRanjbar, Kolja N Eckermann, Hassan M M Ahmed, Héctor M Sánchez C, Stefan Dippel, John M Marshall, and Ernst A Wimmer. Consequences of resistance evolution in a cas9-based sex conversion-suppression gene drive for insect pest management. Proc. Natl. Acad. Sci. U. S. A., 115(24):6189–6194, June 2018.

30. Philip J Lester, Mariana Bulgarella, James W Baty, Peter K Dearden, Joseph Guhlin, and John M Kean. The potential for a CRISPR gene drive to eradicate or suppress globally invasive social wasps. Sci. Rep., 10(1):12398, July 2020.

31. Andrew Hammond, Roberto Galizi, Kyros Kyrou, Alekos Simoni, Carla Siniscalchi, Dimitris Katsanos, Matthew Gribble, Dean Baker, Eric Marois, Steven Russell, et al. A crispr-cas9 gene drive system targeting female reproduction in the malaria mosquito vector anopheles gambiae. Nature biotechnology, 34(1):78–83, 2016.

32. Nikolay P Kandul, Junru Liu, Jared B Bennett, John M Marshall, and Omar Akbari. A home and rescue gene drive efficiently spreads and persists in populations. bioRxiv, 2020.

33. Gerard Terradas, Anna B Buchman, Jared B Bennett, Isaiah Shriner, John M Marshall, Omar S Akbari, and Ethan Bier. Inherently confinable split-drive systems in drosophila. Nature communications, 12(1):1–12, 2021.

34. Nicky R Faber, Gus R McFarlane, R Chris Gaynor, Ivan Pocrnic, C Bruce A Whitelaw, and Gregor Gorjanc. Novel combination of crispr-based gene drives eliminates resistance and localises spread. Scientific reports, 11(1):1–15, 2021.

35. Noble I Egekwu, Francisco Posada, Daniel E Sonenshine, and Steven Cook. Using an in vitro system for maintaining varroa destructor mites on apis mellifera pupae as hosts: studies of mite longevity and feeding behavior. Exp. Appl. Acarol., 74(3):301–315, March 2018.

36. Cameron J Jack, Ping-Li Dai, Edzard van Santen, and James D Ellis. Comparing four methods of rearing varroa destructor in vitro. Exp. Appl. Acarol., 80(4):463–476, April 2020.

37. Maeva A Techer, Rahul V Rane, Miguel L Grau, John M K Roberts, Shawn T Sullivan, Ivan Liachko, Anna K Childers, Jay D Evans, and Alexander S Mikheyev. Divergent evolutionary trajectories following speciation in two ectoparasitic honey bee mites. Communications Biology, 2(1):357, October 2019.

38. Zachary Y Huang, Guowu Bian, Zhiyong Xi, and Xianbing Xie. Genes important for survival or reproduction in varroa destructor identified by RNAi. Insect Sci., 26(1):68–75, February 2019.

39. Wannes Dermauw, Wim Jonckheere, Maria Riga, Ioannis Livadaras, John Vontas, and Thomas Van Leeuwen. Targeted mutagenesis using CRISPR-Cas9 in the chelicerate herbivore tetranychus urticae. Insect Biochem. Mol. Biol., 120:103347, May 2020.

40. Anthony A James. Gene drive systems in mosquitoes: rules of the road. Trends Parasitol., 21(2):64–67, February 2005.

41. Steven P Sinkins and Fred Gould. Gene drive systems for insect disease vectors. Nat. Rev. Genet., 7(6):427–435, June 2006.

42. Thomas A A Prowse, Phillip Cassey, Joshua V Ross, Chandran Pfitzner, Talia A Wittmann, and Paul Thomas. Dodging silver bullets: good CRISPR gene-drive design is critical for eradicating exotic vertebrates. Proc. Biol. Sci., 284(1860), August 2017.

43. Robert L Unckless, Andrew G Clark, and Philipp W Messer. Evolution of resistance against crispr/cas9 gene drive. Genetics, 205(2):827–841, 2017.

44. Charleston Noble, Ben Adlam, George M Church, Kevin M Esvelt, and Martin A Nowak. Current CRISPR gene drive systems are likely to be highly invasive in wild populations. Elife, 7, June 2018.

45. James J Bull. Lethal gene drive selects inbreeding. Evol Med Public Health, 2017(1):1–16, December 2016.

46. Jun Li, Ofer Aidlin Harari, Anna-louise Doss, Linda L Walling, Peter W Atkinson, Shai Morin, and Bruce E Tabashnik. Can CRISPR gene drive work in pest and beneficial haplodiploid species? Evol. Appl., 9:1759, June 2020.

47. Ingemar Fries, Scott Camazine, and James Sneyd. Population dynamics of varroa jacobsoni: A model and a review. null, 75(1):5–28, January 1994.

48. Johan N M Calis, Ingemar Fries, and Stephen C Ryrie. Population modelling of varroa jacobsoni oud. Apidologie, 30(2-3):111–124, 1999.

49. Stephen Martin. A population model for the ectoparasitic mite varroa jacobsoni in honey bee (apis mellifera) colonies. Ecol. Modell., 109(3):267–281, June 1998.

50. Lilia I De Guzman, Thomas E Rinderer, and Amanda M Frake. Growth of varroa destructor (acari: Varroidae) populations in russian honey bee (hymenoptera: Apidae) colonies. Annals ofthe Entomological Society of America, 100(2):187–195, 2007.

51. Alexis L Beaurepaire, Klemens J Krieger, and Robin F A Moritz. Seasonal cycle of inbreeding and recombination of the parasitic mite varroa destructor in honeybee colonies and its implications for the selection of acaricide resistance. Infect. Genet. Evol., 50:49–54, June 2017.

52. Tom J de Jong. Gene drives do not always increase in frequency: from genetic models to risk assessment. Journal of Consumer Protection and Food Safety, 12(4):299–307, 2017.

53. P; Currie Gatien. Timing of acaracide treatments for control of low-level populations of varroa destructor (acari: Varroidae) and implications for colony performance of honey bees. Ottawa Law Rev., 135(5):749–763, October 2003.

54. Willem J Boot, David JA Sisselaar, Johan NM Calis, and Joop Beetsma. Factors affecting invasion of varroa jacobsoni (acari: Varroidae) into honeybee, apis mellifera (hymenoptera: Apidae), brood cells. Bulletin of Entomological Research, 84(1):3–10, 1994.

55. Noble I Egekwu, Francisco Posada, Daniel E Sonenshine, and Steven Cook. Using an in vitro system for maintaining varroa destructor mites on apis mellifera pupae as hosts: studies of mite longevity and feeding behavior. Exp. Appl. Acarol., 74(3):301–315, 2018.

56. Nonno Hasegawa, Maeva Techer, and Alexander S Mikheyev. A toolkit for studying varroa genomics and transcriptomics: preservation, extraction, and sequencing library preparation. BMC Genomics, 22(1):54, January 2021.

57. Luis Medina Medina, Stephen J Martin, Laura Espinosa-Montaño, and Francis L W Ratnieks. Reproduction of varroa destructor in worker brood of africanized honey bees (apis mellifera). Exp. Appl. Acarol., 27(1-2):79–88, 2002.

58. Gloria DeGrandi-Hoffman and Robert Curry. A mathematical model of varroa mite (varroa destructor anderson and trueman) and honeybee (apis mellifera l.) population dynamics. ìni. J. Acarology, 30(3):259–274, September 2004.

59. MD Ifantidis. Parameters of the population dynamics of the varroa mite on honeybees. Journal of Apicultural Research, 23(4):227–233, 1984.

60. L E Brettell and S J Martin. Oldest varroa tolerant honey bee population provides insight into the origins ofthe global decline of honey bees. Sci. Rep., 7:45953, April 2017.

61. Sandra Barroso-Arévalo, Eduardo Fernández-Carrión, Joaquín Goyache, Fernando Molero, Francisco Puerta, and José Manuel Sánchez-Vizcaíno. High load of deformed wing virus and varroa destructor infestation are related to weakness of honey bee colonies in southern spain. Front. Microbiol., 10:1331, June 2019.

62. Gennaro Di Prisco, Desiderato Annoscia, Marina Margiotta, Rosalba Ferrara, Paola Varricchio, Virginia Zanni, Emilio Caprio, Francesco Nazzi, and Francesco Pennacchio. A mutualistic symbiosis between a parasitic mite and a pathogenic virus undermines honey bee immunity and health. Proc. Natl. Acad. Sci. U. S. A., 113(12):3203–3208, March 2016.

63. R W Currie and P Gatien. Timing acaricide treatments to prevent varroa destructor (acari: Varroidae) from causing economic damage to honey bee colonies. Canadian Entomologist; Ottawa, 138(2):238–252, April 2006.

64. Keith S Delaplane and W Michael Hood. Economic threshold for varroa jacobsoni oud. in the southeastern USA. Apidologie, 30(5):383–395, 1999.

65. R M Goodwin, M A Taylor, H M Mcbrydie, and H M Cox. Drift of varroa destructor-infested worker honey bees to neighbouring colonies. J. Apic. Res., 45(3):155–156, January 2006.

66. David T Peck, Michael L Smith, and Thomas D Seeley. Varroa destructor mites can nimbly climb from flowers onto foraging honey bees. PLoS One, 11(12):e0167798, December 2016.

67. David Thomas Peck and Thomas Dyer Seeley. Mite bombs or robber lures? the roles of drifting and robbing in varroa destructor transmission from collapsing honey bee colonies to their neighbors. PLoS One, 14(6):e0218392, June 2019.

68. Thomas D Seeley and Michael L Smith. Crowding honeybee colonies in apiaries can increase their vulnerability to the deadly ectoparasite varroa destructor. Apidologie, 46(6): 716–727, November 2015.

69. Sean P Leonard, J Elijah Powell, Jiri Perutka, Peng Geng, Luke C Heckmann, Richard D Horak, Bryan W Davies, Andrew D Ellington, Jeffrey E Barrick, and Nancy A Moran. Engineered symbionts activate honey bee immunity and limit pathogens. Science, 367(6477): 573–576, 2020.

70. R Core Team et al. R: A language and environment for statistical computing, 2013.

71. R Chris Gaynor, Gregor Gorjanc, and John M Hickey. Alphasimr: An r-package for breeding program simulations. BioRxiv, 2020.

72. M Delia Allen. The effect of a plentiful supply of drone comb on colonies of honeybees. Journal of apicultural research, 4(2):109–119, 1965.

73. R Büchler, A Uzunov, M Kovačić, J Prešern, M Pietropaoli, F Hatjina, B Pavlov, L Charistos, G Formato, E Galarza, et al. Summer brood interruption as integrated management strategy for effective varroa control in europe. Journal of Apicultural Research, pages 1–10, 2020.

74. SJ Martin and D Kemp. Average number of reproductive cycles performed by varroa jacobsoni in honey bee (apis mellifera) colonies. Journal of Apicultural research, 36(3-4): 113–123, 1997.

75. Ingemar Fries and Peter Rosenkranz. Number of reproductive cycles of varroa jacobsoni in honey-bee (apis mellifera) colonies. Experimental & applied acarology, 20(2):103–112, 1996.

76. S Fuchs. Choice in varroa jacobsoni oud. between honey bee drone or workerbrood cells for reproduction. Behavioral Ecology and Sociobiology, 31(6):429–435, 1992.

77. Peter Rosenkranz, Pia Aumeier, and Bettina Ziegelmann. Biology and control of varroa destructor. Journal of invertebrate pathology, 103:S96–S119, 2010.

78. Francesco Nazzi and Yves Le Conte. Ecology of varroa destructor, the major ectoparasite of the western honey bee, apis mellifera. Annu. Rev. Entomol., 61: 417–432, 2016.

79. S J Martin. Reproduction of varroa jacobsoni in cells of apis mellifera containing one or more mother mites and the distribution of these cells. J. Apic. Res., 34(4):187–196, January 1995.

